# HS-AFM Reveals Hierarchical RNA Folding Driving Condensate Assembly and Material States

**DOI:** 10.64898/2025.12.08.693082

**Authors:** S M Neaz Mahmud, Kenichi Umeda, Tamoghna Das, Nadia Shoukat, Noriyuki Kodera, Hanae Sato

## Abstract

Liquid-liquid phase separation (LLPS) is a fundamental mechanism of intracellular biomolecular condensation formation. While RNA self-assembly has been implicated in condensate formation, it remains unclear how RNA condensation translates into distinct physical behaviors at the nanoscale. Utilizing HS-AFM, this study characterizes RNA condensation dynamics with single-molecule resolution. Imaging captured the transition from individual RNA folding to intermolecular clustering, ultimately leading to progressive condensate assembly. Beyond morphological description, condensate behavior was further examined through fusion dynamics and mechanical response. Post-fusion shape evolution quantified how merged condensates recover circular morphology over time, providing a dynamic readout of material behavior. Nanomechanical properties were independently assessed through force-curve measurements. Together, these analyses consistently distinguish the droplet-like poly A condensates from fractal-shaped assemblies of total RNA. Taking together, these findings directly link RNA folding dynamics, condensate assembly, and emergent physical properties, establishing a quantitative framework for defining condensate material states at the nanoscale.

## Introduction

Liquid-liquid phase separation (LLPS) is a physiological process that separates homogenous solutions into two or more coexisting phases. It occurs when multiple components of a homogenous solution separate into distinct liquid phases with varying concentrations^[1,2]^. LLPS is influenced by various factors, including external environment such as temperature, pressure and pH as well as the introduction of energy into the molecular environment, and alterations in component properties like polarity, hydrophobicity and hydrophilicity. In cells, LLPS generates liquid droplet-like structures that form dynamic, membraneless biomolecular condensates capable of rapidly exchanging molecules with their surrounding^[3,4]^. These condensates coordinate cellular organization and biochemical activity, highlighting LLPS as a central mechanism for the formation and functional regulation of cellular activities^[5]^. Multiple interactions among biomolecules can drive their physiological transition from one phase to another^[6,7]^. These condensates typically exhibit higher density and slower molecular motion compared to the surrounding medium, which facilitates elevated rates of biochemical reactions^[5,6,8]^. Mostly, these structures exhibit liquid characteristics and are commonly described as puncta, droplets and condensates^[9]^, exemplified by transcriptional condensates, nucleoli, stress granules (SGs), processing bodies (P-bodies), Cajal bodies, and many other membraneless organelles^[10,11]^. Biomolecular condensates are composed of protein and RNA molecules where proteins solely or in combination with RNA form granules^[12,13]^. Importantly, RNA alone can undergo condensation in vitro without proteins, demonstrating that its fundamental properties are sufficient to initiate LLPS^[14]^. Intermolecular RNA interactions are thought to primarily drive RNA only phase separation under appropriate physiological conditions. However, the role of RNA-RNA interaction in biomolecular condensation is less understood compared to RNA-protein interactions in granule formation^[15,16]^.

In vitro RNA condensation is promoted by high concentrations of homotypic or heterotypic RNA molecules along with polyvalent cations (e.g. cobalt hexamine, spermine, poly-lysine, Ca^2+^, Mg^2+^) and/or crowding agents such as polyethylene glycol (PEG)^[17–19]^. In the presence of divalent cations, GC-rich RNAs with tripeptide repeats can transition from liquid to gel in vitro, suggesting a potential mechanism for the formation of intracellular RNA granules through LLPS in RNA repeat expansion disorders^[19,20]^. Similarly, RNA self-assembly is predominantly driven by robust and redundant RNA-RNA interactions, highlighting its role in LLPS^[14]^. At elevated concentrations, RNA molecules can form intermolecular interactions when in close spatial proximity, particularly for RNAs with highly flexible secondary structures^[21]^. Weak, transient, multivalent interactions, including π-π and π-cation interactions, participate in the formation of RNA-driven LLPS ^[22,23]^. These observations underscore the importance of intrinsic RNA properties in LLPS formation, as intermolecular interactions depend on the RNA structure and sequence composition. The three-dimensional structure of RNA determines its biological activities, making the studies of RNA structure essential for understanding RNA sequence and structural features^[24]^.

Atomic force microscopy (AFM) utilizes the interaction forces between the end of a probe tip and the sample surface, enabling nanometer-resolution imaging of biomolecules^[25,26]^. In particular, high-speed AFM (HS-AFM) offers high spatiotemporal resolution, allowing real-time visualization of the dynamic behavior of biomolecules such as DNA, RNA, and proteins^[27–29]^. In this study, we used HS-AFM to track the real-time, single-molecule dynamics of RNA-driven LLPS, examining how intra and intermolecular RNA interactions govern phase separation. This approach enables high-resolution detection of LLPS at the single-molecule level, overcoming the limitations of conventional optical imaging, which typically cannot detect single molecules and only visualize larger droplets formed at high RNA concentrations. Measurement of fusion kinetics, Young’s modulus and indentation depth confirms the mechanical properties of RNA condensation with significant differences among poly A droplets and total RNA aggregates. Our method also allows us to capture the full transition of RNA-driven LLPS from single molecules to condensates, showing that even a few molecules can form droplets, and that LLPS proceeds in a stepwise manner from single molecule RNA folding mediated by cis interactions, through intermolecular interactions, to the formation of higher order assemblies.

## Results

### Effects of Crowding Agent and Ionic Conditions on RNA-driven LLPS

To investigate the mechanism of RNA-driven LLPS, we used two types of RNAs widely employed in previous studies: in vitro synthesized poly A RNA, a simple homopolymeric model, and total RNA isolated from HEK293T cell line, representing the complexity and diversity of the cellular transcriptome. First, we tested the condensation formation of selected RNAs in the presence of PEG, a commonly used crowding agent that promotes LLPS by enhancing biomolecular interactions and stabilizing droplets in vitro, to assess their phase separation capability under bright-field microscopy^[30]^. Under the condition of 10% PEG, 10 mM MgCl₂, and 10 mM Tris-HCl, poly A RNA (150 ng/µL) formed spherical, liquid-like droplets, whereas total RNA (150 ng/µL) predominantly formed aggregates, suggesting different condensation behavior as observed in bright-field microscopy (Figure S1a). Consistent with previous studies, total RNA transitioned into more circular, droplet-like structures resembling those formed by poly A upon heat incubation, suggesting that thermal fluctuations can overcome structural or sequence-based barriers to liquid-like phase separation (Figure S1a)^29,30^.

Although PEG-based buffers are commonly used to induce RNA-driven LLPS in vitro, they require high RNA concentrations (>150 ng/µL), leading to a densely condensed molecular state in which HS-AFM can no longer resolve individual RNA structures. This poses a technical challenge, as LLPS typically requires high RNA concentrations, whereas HS-AFM requires much lower concentrations to resolve molecular transitions due to its scanning-based detection. To overcome this limitation, we established an alternative approach using elevated Mg^2+^ concentration at lower RNA concentrations, as previously reported for RNA-driven LLPS in the absence of crowding agents ^[31]^. This strategy enables RNA condensation while maintaining single-molecule resolution. Mg^2+^ primarily neutralizes the negative charges of RNA, thereby reducing electrostatic repulsion between RNA strands and within folded RNA structures. In addition, it promotes the formation of RNA tertiary structures such as hairpins, pseudoknots, and long-range interactions^[32]^. By electrostatically binding to the phosphate backbone, Mg^2+^ also facilitates π–π stacking, base pairing, and hydrogen bonding among or within RNA strands, highlighting its significant structural and functional roles in RNA behavior^[33]^.

First, we confirmed RNA condensation at 100 mM Mg^2+^ without PEG at the same RNA concentration (150 ng/µL). Bright-field imaging showed that poly A RNA (150 ng/µL) formed circular droplets, whereas total RNA (150 ng/µL) produced irregular aggregates, similar to those observed in PEG-containing buffers (Figure 1a).^34^

**Figure 1.**
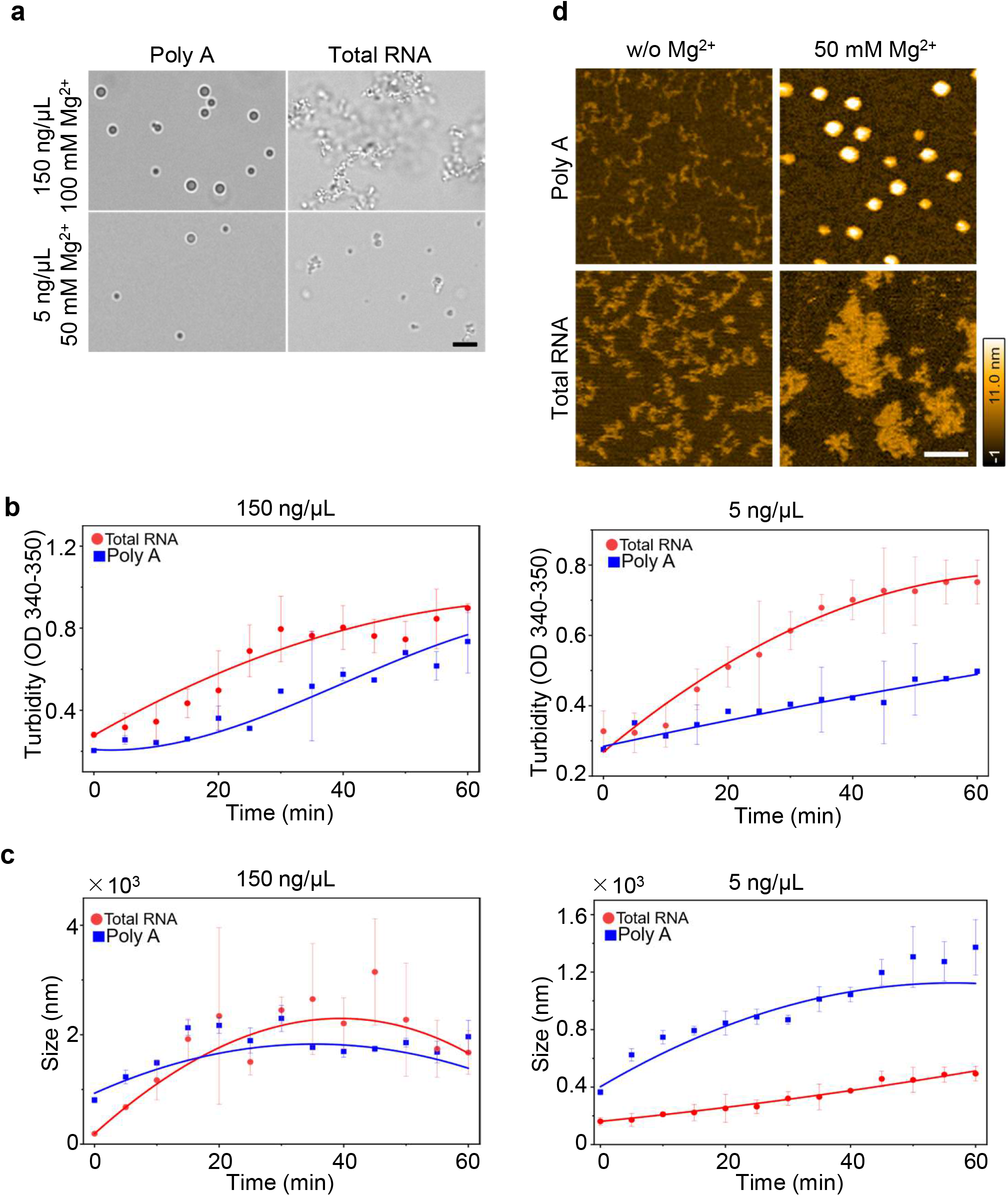
From bulk to single molecules: HS-AFM reveals RNA condensation at low concentrations. a. Brightfield microscopic images showing RNA condensates of poly A and total RNA. 150 ng/µL RNA with 100 mM Mg2+ and 5 ng/µL RNA with 50 mM Mg2+ was incubated at room temperature for 10–15 minutes before imaging (Scale bar 10 µm). b. Turbidity at 340-350 nm (OD 340-350) was measured for poly A and total RNA condensates where 150 ng/µL of RNA with 100 mM Mg2+ (left panel) and 5 ng/µL RNA with 50 mM Mg2+ (right panel) of both poly A and total RNA were used. The solid line shows mean ± SD of measurement done for three independent samples prepared under same condition. c. The size (diameter) evolution of droplets was followed by DLS in experiment condition (b). The solid line shows mean ± SD of measurement done for three independent samples prepared under same condition. d. HS-AFM images of RNA condensations formed by poly A and total RNA at 5 ng/µL RNAs with 50 mM Mg2+. Experiments were conducted within a 350 nm x 350 nm scanning area with 120 x 120 pixels (Scale bar 100 nm).

After confirming that elevated Mg²⁺ in PEG-free buffer induces RNA condensation, RNA concentrations and buffer conditions were optimized for HS-AFM imaging. A minimum of 25 mM Mg²⁺ and RNA concentrations as low as 5 ng/µL were sufficient to induce condensation in the absence of PEG (Figure S1b and 1a). Although condensates remained below the sensitivity threshold of bright-field microscopy at these concentrations (Figure 1a, 5 ng/ul), HS-AFM enabled sensitive visualization of early-stage RNA condensation.

Turbidity at 340–350 nm was measured to monitor bulk condensation, reflecting overall light scattering from RNA assemblies. Turbidity increased over time for both total RNA and poly A at 5 and 150 ng/µL, with a more rapid and pronounced increase observed for total RNA at both concentrations (Figure 1b). Because turbidity depends not only on particle size but also on particle number and structural heterogeneity, the stronger signal observed for total RNA likely reflects the formation of a larger number of heterogeneous light-scattering assemblies. In contrast, poly A exhibited consistently lower turbidity, suggesting fewer and/or more uniform condensates despite measurable particle growth.

Dynamic light scattering (DLS) further characterized particle size distributions. At low concentration (5 ng/µL), poly A formed larger average particles than total RNA, whereas total RNA remained smaller but more broadly distributed (Figure 1c, right panel). At higher concentration (150 ng/µL), both RNA types exhibited time-dependent growth that eventually reached a plateau, resulting in similar average particle sizes (Figure 1c, left panel). For poly A, assemblies initially measured approximately 0.3×10^3^ nm in diameter and gradually increased to ∼1.5×10^3^ nm over ∼60 minutes (Figure 1c, left panel). This relatively uniform growth was reflected in a low polydispersity index (PDI = 0.33), consistent with progressive coarsening behavior characteristic of Ostwald ripening (Figure S1c, left panel) ^[34]^. In contrast, total RNA assemblies exhibited multimodal size distributions (PDI = 0.69) (Figure S1c, left panel), indicating heterogeneous assembly rather than uniform coarsening. Similar PDI values were also observed for lower concentration (5 ng/µL) of both RNAs (Figure S1c, right panel). Together, these results indicate that lower concentration of RNA is capable of condensation formation in suitable condition and the average particle size alone does not account for bulk turbidity differences. Instead, sequence and structural diversity in total RNA likely promote heterogeneous and multivalent intermolecular interactions, generating irregular clusters that enhance light scattering. In contrast, the homopolymeric nature of poly A favors more uniform assembly pathways, resulting in lower turbidity despite comparable or even larger particle sizes under certain conditions.

Using these lower RNA concentrations, HS-AFM imaging revealed that poly A formed fine droplet-like structures, whereas total RNA formed irregular aggregated assemblies in 50 mM Mg²⁺ (Figure 1d). In the absence of Mg²⁺, only thread-like RNA structures were observed, indicating limited intermolecular interactions (Figure 1d). The concordance between real-time single-molecule imaging and bulk DLS measurements indicates that the observed condensation behavior is consistent across both surface-based and solution-phase analyses. Together, these results demonstrate that RNA condensation can be induced under defined buffer conditions and that poly A and total RNA exhibit distinct assembly behaviors detectable at both bulk and single-molecule levels.

### Real-time visualization of poly A RNA condensation Dynamics

We next aimed to capture the dynamic process of poly A RNA-driven LLPS, monitoring the progression of condensates from linear poly A RNA molecules to fully formed droplets. Conventional bright-field microscopy typically requires defined incubation periods prior to observation and lacks the spatial resolution to detect early nanoscale intermediates, making the initial stages of condensation difficult to capture. In contrast, HS-AFM enables real-time imaging at single-molecule resolution under near-physiological conditions, allowing direct observation of early assembly events and condensate maturation dynamics.

At a concentration of 5 ng/µL, linear poly A RNA was observed at room temperature on a mica surface for long time (600 s). Upon addition of 50 mM Mg^2+^, linear poly A RNA underwent partial folding followed by droplet formation, capturing the ongoing transition from folded RNA to droplets (Figure 2a & Movie 1).

**Figure 2.**
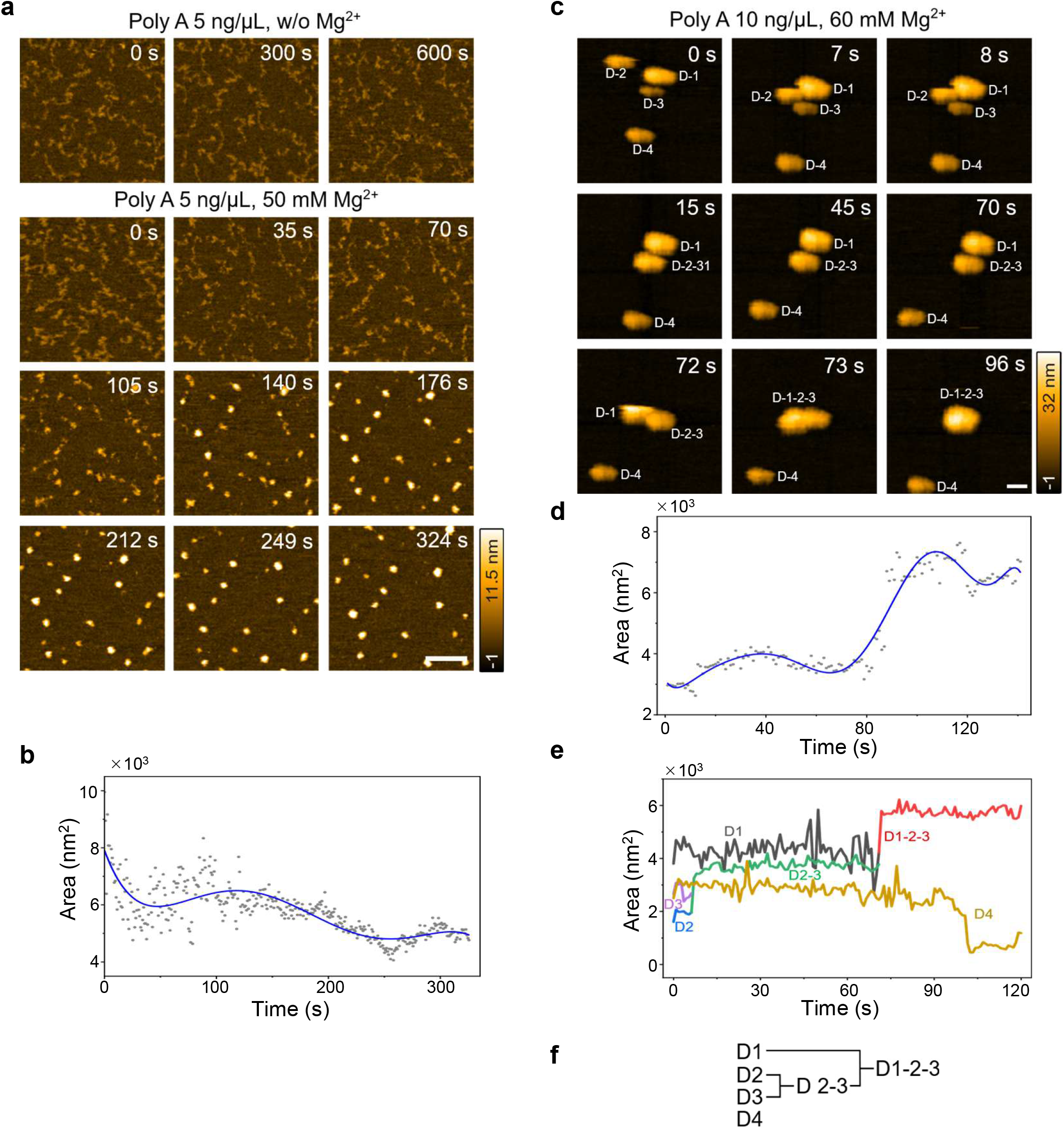
HS-AFM observation of droplet-like progression of poly A RNA condensation. a. Successive HS-AFM images showing real-time droplet formation by poly A RNA upon addition of Mg²⁺. Poly A RNA (5 ng/µL) and 50 mM Mg²⁺ were applied to a mica surface at room temperature. Imaging was performed in a 500 × 500 nm scanning area with 120 × 120 pixels (Scale bar, 100 nm). b. Scatter plots showing changes in droplet area under the conditions in (a). Total area decreases over time due to RNA folding and condensation. The solid line represents a polynomial fit to the experiment data, illustrating the decreasing trend of area during droplet formation. c. HS-AFM frames showing droplet maturation at higher RNA and Mg²⁺ concentrations. Poly A RNA (10 ng/µL) and 60 mM Mg²⁺ were used to capture fusion events. Imaging was performed in a 350 × 350 nm scanning area with 120 × 120 pixels (Scale bar, 50 nm). d. Scatter plots showing changes in area during droplet maturation under the conditions in (c). Area increases over time as droplets fuse or merge and form large droplets. The solid line represents a polynomial fit to the experiment data, illustrating the increasing trend of area during droplet fusion. e. Line graph showing the dynamics of area change during maturation of droplets. At first, two droplets (D2 and D3) merge and form droplet (D-2-3) which finally merged with (D-1) and form droplet (D-1-2-3). f. Schematic illustrating the progression of droplet maturation under the condition in (c).

As poly A is a homopolymer lacking complementary sequences, its compaction under cationic conditions arises without canonical base-pairing interactions. Instead, Mg²⁺ mediated charge screening likely promotes base-stacking interactions and intramolecular collapse. This compaction may facilitate subsequent intermolecular association, ultimately leading to droplet formation. To quantify the structural changes of linear poly A RNA during droplet formation, we measured the area, height, and circularity of RNA molecules over time (Figure 2b & S2a, S2b). During this process, the area of RNA molecules decreased from ∼7000 nm^2^ to ∼4000 nm^2^ (Figure 2b), the height increased from ∼3 nm to over 8 nm (Figure S2a), and circularity increased from ∼0.3 to ∼0.9 (Figure S2b), indicating progressive compaction and droplet formation. The average size of droplets observed by HS-AFM after 5 minutes of Mg^2+^ addition was approximately 582 nm², which is comparable in scale to the particle size measured by DLS for 5 ng/µL poly A RNA at the same time point (Figure S2c & S2d), supporting consistency between single-molecule imaging and bulk measurements. At 10 ng/µL RNA and 60 mM Mg^2+^, we observed droplet fusion, in which neighboring small droplets coalesced into larger condensates (Figure 2c & Movie 2). Measurements of area, height, and circularity of individual droplets showed increases over time due to droplet merging (Figure 2d, S2e & S2f). To characterize droplet maturation, individual condensates were tracked in real time by assigning unique particle IDs and monitoring their projected area over time (Figure 2e,f). This approach enabled direct tracing of droplet lineage, allowing us to determine which specific droplets merged and how individual assemblies contributed to the growth of larger condensates. Such real-time tracking provides mechanistic insight into the fate of individual molecular assemblies during coalescence, revealing stepwise fusion events that drive droplet maturation. Collectively, these results suggest a two-step condensation mechanism: (i) intramolecular folding and compaction, leading to reduced molecular footprint, followed by (ii) intermolecular droplet maturation via fusion, which drives expansion of condensate size and morphological optimization. Importantly, HS-AFM captures the entire process of droplet formation in real time, from linear RNA to single-stranded folding, formation of single-molecule droplets, and finally to mature droplets, further revealing the molecular transitions of RNA condensation.

### Real-time visualization of Total RNA Condensation Dynamics

Condensation formed by total RNA is of particular interest, as it may more closely reflect the mechanisms underlying ribonucleoprotein (RNP) granule formation in cells. Several studies have suggested that such condensates are driven by intermolecular RNA–RNA base-pairing interactions^29^, which have been predicted through RNA secondary structure modeling but have not yet been directly visualized. While the system supports heating up to 70 °C, imaging at temperatures exceeding 90 °C, which induces droplet-like structures, is not compatible with HS-AFM imaging. Therefore, imaging was performed under non-heated conditions to enable real-time observation. At 5 ng/µL, total RNA was observed as non-linear, pre-folded structures on the mica surface, exhibiting limited local fluctuations and no substantial intermolecular interactions in the absence of Mg^2+^. Upon addition of 50 mM Mg^2+^, total RNA became dynamically mobile and began interacting with other RNA molecules, eventually forming condensate clusters that appeared as more compact, tightly packed structures (Figure 3a, movie 3). We quantified the area, height and circularity throughout the imaging period to assess morphological changes. Following Mg^2+^ addition, the height and circularity increased (Figure S3a,b), while the area fluctuated with repeated increases and decreases (Figure 3b). These dynamics suggest that structured total RNA first assembled (∼150 s), followed by intermolecular RNA–RNA interactions that trigger molecular compaction during condensate maturation. Comparison of aggregate size at 5 minutes between HS-AFM and DLS measurements revealed similar behavior. Small differences in absolute size likely reflect detection sensitivity, as very short RNA species detectable by DLS may be underrepresented in HS-AFM imaging (Figure S2c & S2d).

**Figure 3.**
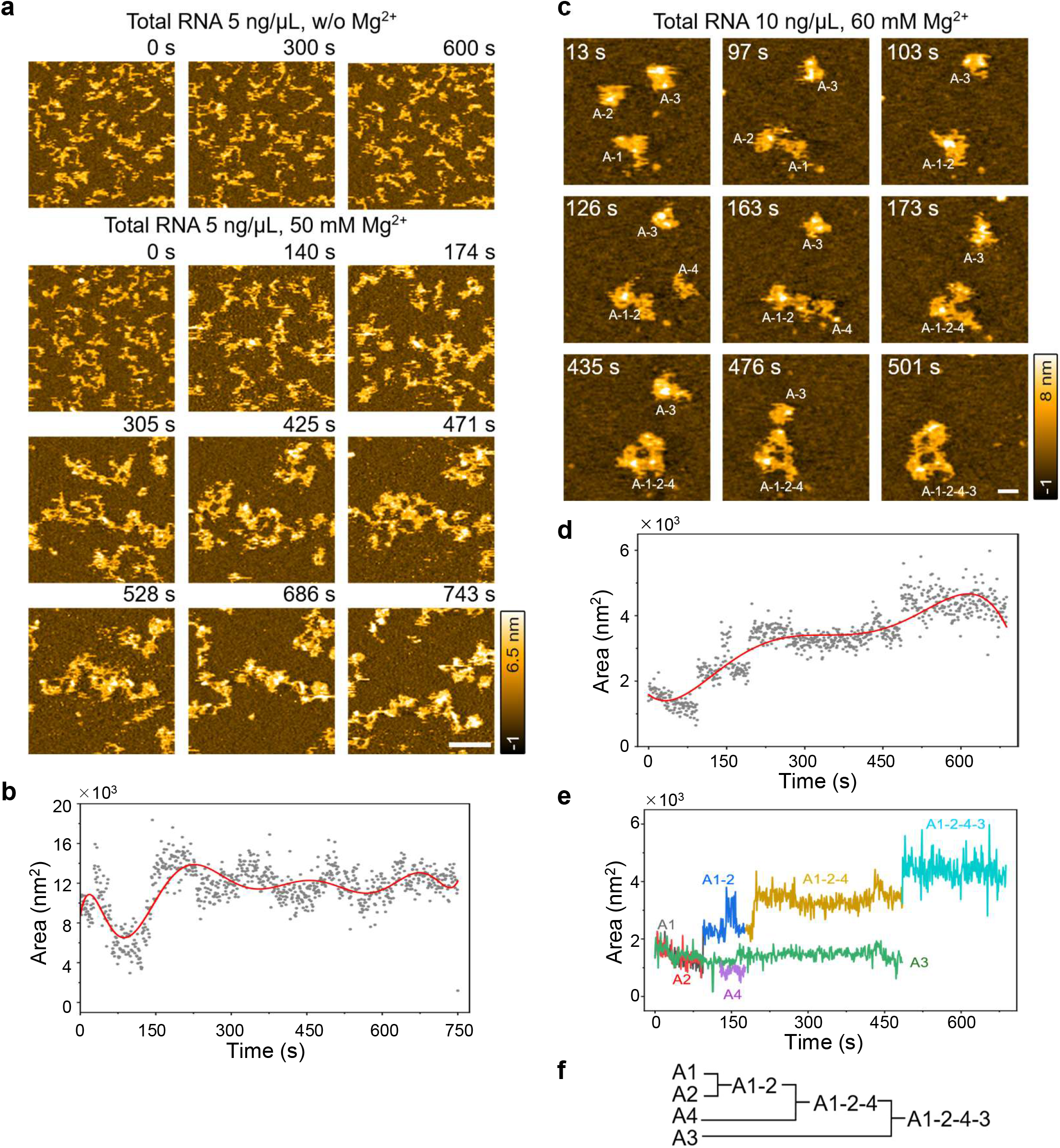
HS-AFM observation of aggregation-like progression of total RNA condensation. a. Successive HS-AFM images showing the effects of Mg²⁺ on total RNA. Total RNA form aggregates upon the addition of 50 mM Mg2+. Imaging was performed in a 500 × 500 nm scanning area with 120 × 120 pixels (Scale bar, 100 nm). b. Scatter plot showing changes in area of total RNA during Mg²⁺ induced aggregation under the conditions in (a). The solid line represents a polynomial fit to the experiment data, illustrating the trend of area during aggregation. c. HS-AFM frames showing the process of large aggregation formation of total RNA at high RNA and Mg²⁺ concentrations (10 ng/µL total RNA, 60 mM Mg²⁺). Imaging was performed in a 350 × 350 nm scanning area with 120 × 120 pixels (Scale bar, 50 nm). d. Scatter plot showing changes in area during aggregate formation under the conditions in (c). The area increases gradually as RNAs accumulate in the growing aggregate. The solid line represents a polynomial fit to the experiment data, illustrating the increasing trend of area during aggregate formation. e. Line graph showing the dynamics of area change during the formation of large aggregates. At the beginning, aggregates A-1 and A-2 interact to form A-1-2 and gradually form A-1-2-4-3 after merging with aggregates A4 and A5 respectively. f. Schematic illustrating the progression of droplet maturation.

To characterize aggregate maturation, individual condensates were tracked in real time by assigning unique particle IDs and monitoring their projected area over time at 10 ng/µL RNA and 60 mM Mg^2+^ (Figure 3c–f), as described for poly A in Figure 2c–f. In contrast to poly A condensation, total RNA aggregates exhibited significant size fluctuations following cluster merging, suggesting dynamic intermolecular structural rearrangements after coalescence. These differences indicate distinct maturation behaviors between poly A and total RNA assemblies. Together, these results support a model in which pre-folded total RNAs undergo an initial intermolecular assembly phase (area increase), followed by Mg^2+^ dependent compaction into condensates. Early RNA–RNA interactions appear to initiate cluster formation, with subsequent structural rearrangements within clusters that ultimately drive compaction.

### Droplet-like vs. Aggregate-like Condensation: Poly A Exhibits Faster Fusion Kinetics than Total RNA

A defining material property of liquid-like condensates is their ability to coalesce and rapidly recover a spherical morphology. The characteristic timescale of this post-coalescence shape recovery, termed as relaxation time (τ), reflects intrinsic material parameters such as viscosity and surface tension ^[35]^. We therefore quantified recovery kinetics to directly compare the material states of poly A and total RNA assemblies.

Circularity was monitored from the onset of coalescence (t = 0) until morphological stabilization (Figure 4a). Poly A condensates reached geometric equilibrium rapidly, plateauing within ∼1.63 s (Figure 4b,d). In contrast, total RNA assemblies exhibited a nonlinear and fluctuating recovery profile, with a significantly longer average τ (∼8.32 s) (Figure 4c,d).

**Figure 4.**
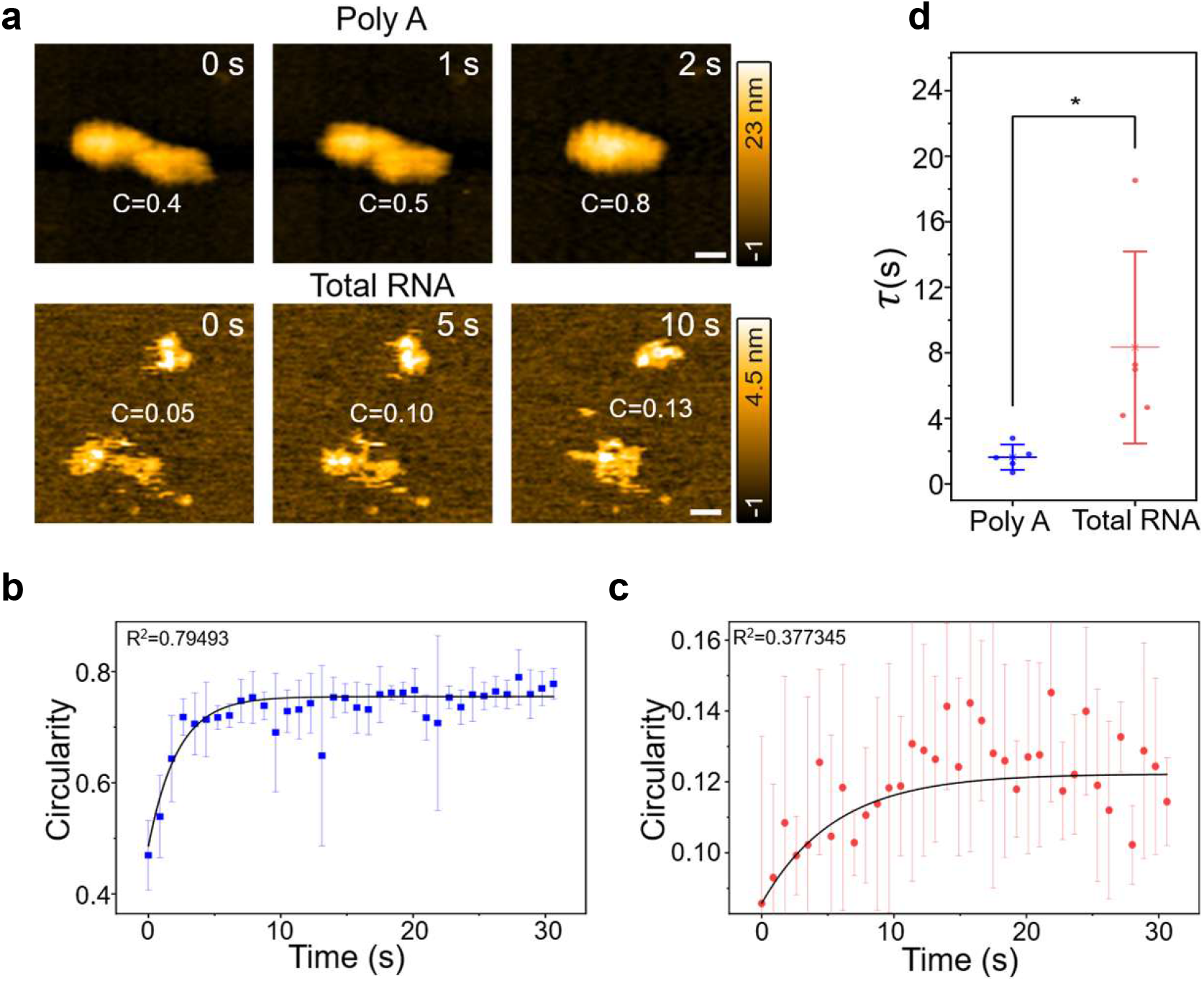
Comparison of fusion relaxation profile of poly A droplets and total RNA aggregates. a. Representative time-lapse topography HS-AFM images depicting the coalescence of poly A and total RNA aggregates. The morphological evolution is characterized by a time-dependent increase in circularity, indicating a transition from asymmetric intermediate structures to a thermodynamically stable spherical state. The plateau in circularity values at the post-fusion stage confirms the completion of the relaxation process (Scale bar, 50 nm). b. Scatter plot representing the morphological relaxation of poly A condensates from the onset of fusion (t = 0s) to 30s post-fusion. Data points represent the mean± SD (n = 5). The solid black line denotes polynomial regression curve, highlighting the kinetics of the transition toward a steady-state spherical geometry. c. Scatter plot showing the evolution of circularity from the initial contact (t = 0s) through 30s of interaction for total RNA aggregates. Data points represent the mean ± SD (n = 5). The thick solid line denotes a polynomial regression curve, illustrating the transition of the total RNA assemblies toward a stabilized post-interaction state. d. Scatter dot plot showing difference in relaxation time (τ) of poly A and total RNA. Bar graphs representing mean ± SD of samples (n = 5; *p < 0.05, **p < 0.01, ***p < 0.001). A paired-sample t-test was performed to assess differences in means at a 0.05 confidence level.

The rapid equilibration of poly A condensates is consistent with fluid-like behavior governed by efficient fusion and capillary-driven rounding. By contrast, the delayed and irregular recovery dynamics of total RNA assemblies indicate structurally constrained, aggregate-like material properties likely stabilized by multivalent RNA–RNA interactions. Thus, real-time extraction of τ provides a quantitative and operational framework to distinguish droplet-like from aggregate-like condensates based on material behavior rather than morphology alone.

### Mechanical property differences between droplet-like and aggregate-like RNA condensates

The mechanical properties of RNA assemblies determine whether they behave as droplet-like (soft, deformable) or aggregate-like (stiff, solid-like) structures. AFM not only visualizes surface structures but also enables mechanical characterization through force curve measurements. To determine the mechanical properties of poly A droplets and total RNA aggregates, we performed 3D force-mapping measurements. As shown in (Figure 5a,b), reconstructed topographic images correspond to a constant-force surface (0.04 nN) extracted from the 3D datasets clearly resolved each structure. In the approach curve of force measurements obtained on poly A, the force gradually increased, briefly decreased, and then increased again. This behavior suggests that the probe penetrates the liquid-like droplet structure. Despite probe penetration into the poly A droplet, the topographic structure could still be reconstructed, indicating a reversible structural response in which the poly A droplet recovers after deformation, consistent with liquid-like mechanical behavior. Similar hysteresis was also observed in total RNA aggregates. Young’s modulus was calculated by fitting each pixel-wise force curve to the Hertz contact model, using the initial rising region as indicated by the purple dashed lines in (Figure 5c,d). The reconstructed modulus maps (Figure 5e,f) showed both poly A droplets and total RNA aggregates were softer relative to the substrate. Note that the non-uniform modulus distribution on the substrate in the aggregate measurements is attributed to the deposition of condensates on the surface.

**Figure 5.**
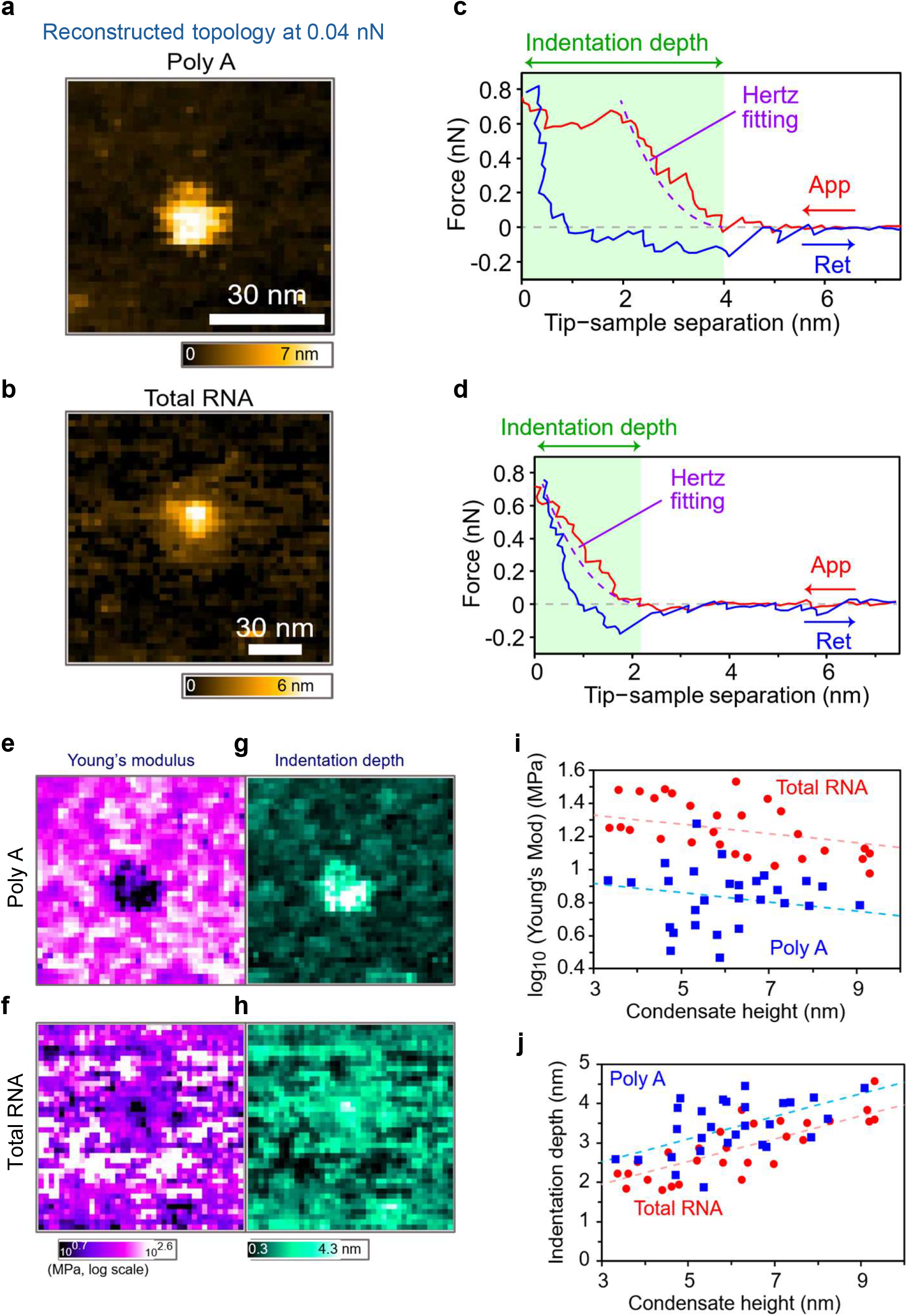
3D force-mapping measurements on RNA condensates. a.–d. Isoforce surface topography images reconstructed from 3D force maps acquired on poly A droplets (a, c) and total RNA aggregates (b, d), together with representative force curves of both condensates (c, d). e. –h. Young’s modulus maps (e, f) and indentation depth maps (g, h) derived from analysis of 3D force maps measured on poly A (e, g) and total RNA aggregates (f, h). i. j. Statistical analysis of dataset-averaged Young’s modulus (i) and indentation depth (j) aggregated across multiple measurements (poly A droplets, n = 28; total RNA aggregates, n = 29), where n denotes the number of analyzed droplets.

To quantify the depth of the probe penetration, we also analyzed the indentation depth, defined as the distance from the onset of force rise to contact with a rigid surface (the green arrows in (Figure 5c,d). As shown in (Figure 5g,h) indentation depth was markedly larger on poly A droplets than on the substrate, with a similar increase observed on total RNA aggregates.

For multiple datasets containing condensates with sizes in the 3–10 nm range, the two metrics were averaged for each dataset and subsequently analyzed statistically. As shown in (Figure 5i), the Young’s modulus of poly A droplets was distributed in the range of approximately 3–10 MPa, whereas total RNA aggregates showed a higher distribution of approximately 10–40 MPa. On average, the modulus of aggregates was about 2.5 times greater than that of poly A droplets. In contrast, indentation depth values for both structures were distributed in the range of approximately 2–4 nm; however, poly A droplets exhibited indentation depths that were on average about 1.25 times greater than those of total RNA aggregates (Figure 5j).

Because both metrics showed a weak dependence on condensation height, we performed an analysis of covariance (ANCOVA) to determine whether statistically significant differences remained after accounting for height effects. The analysis yielded p-values of 4.857 × 10^-13^ for Young’s modulus and 1.343 × 10^-4^ for indentation depth, indicating statistically significant differences in both cases. These findings suggest that poly A droplets are mechanically softer and more readily penetrated by the probe than total RNA aggregates, consistent with their liquid-like physical character.

### Single-molecule investigation of poly A and total RNA condensation

Following characterization of fusion behavior, single-molecule observation was performed to resolve the molecular events underlying RNA condensation, providing real-time insights into RNA folding and droplet assembly at the level of individual molecules. This approach enables tracking of how individual RNA strands fold, interact, and coalesce into larger liquid-like assemblies, revealing the molecular mechanisms driving droplet formation. To achieve clear single-molecule imaging, 0.3 ng/µL poly A RNA was used. Upon addition of 25 mM Mg^2+^ to the scanning chamber during imaging, we observed structural transitions of poly A RNA (Figure 6a; Movie 4). Addition of Mg^2+^ induced intra-molecular (*cis*) interactions, leading to RNA folding and formation of globular shape. Analysis of area and length revealed a reduction in both spatial extent and length of the RNA molecules (Figure S4a & S4b), while the height and circularity increased from 1.8 nm to around 4 nm and from ∼0.1 to ∼0.4 over time, respectively (Figure S4c & S4d), indicating compaction of RNAs induced by Mg^2+^. Although thread-like, unfolded RNAs were rarely observed at a total RNA concentration of 5 ng/µL (Figure 3a), we detected single, thread-like total RNA molecules at lower RNA concentrations (0.5 ng/µL) in the presence of Mg^2+^. HS-AFM observations visualized no major structural transitions of thread-like RNA structure on mica surface in scanning buffer (without Mg^2+^) for long periods of time (Figure 6b). Addition of 25 mM Mg^2+^ to the scanning chamber during scanning triggered RNA folding into a globular shape through *cis* interactions (Figure 6b & Movie 5). Intermolecular (RNA-RNA) interaction then emerged even between closely positioned molecules, highlighting the role of RNA secondary structure in condensate formation (Movie 5). Quantitative analysis showed that area and length decreased (Figure S4e & S4f), while height and circularity increased over time (Figure S4g &S4h), indicating compaction of RNAs induced by Mg^2+^ through self-folding. Poly A RNA compacted rapidly and smoothly, consistent with isotropic base-stacking and low surface tension anisotropy, whereas total RNA folded slowly and irregularly, forming non-spherical assemblies with heterogeneous morphology and limited coalescence. These results reveal distinct cis-interaction behaviors between a homopolymeric RNA (poly A) and a heterogeneous mixture of RNA species (total RNA). The uniform backbone and base-stacking interactions of poly A RNA likely enable isotropic molecular interactions and low surface tension anisotropy, while the sequence heterogeneity and structured regions of total RNA result in irregular, non-spherical, and sometimes gel-like aggregates, reflecting kinetically trapped, non-equilibrium assemblies.

**Figure 6.**
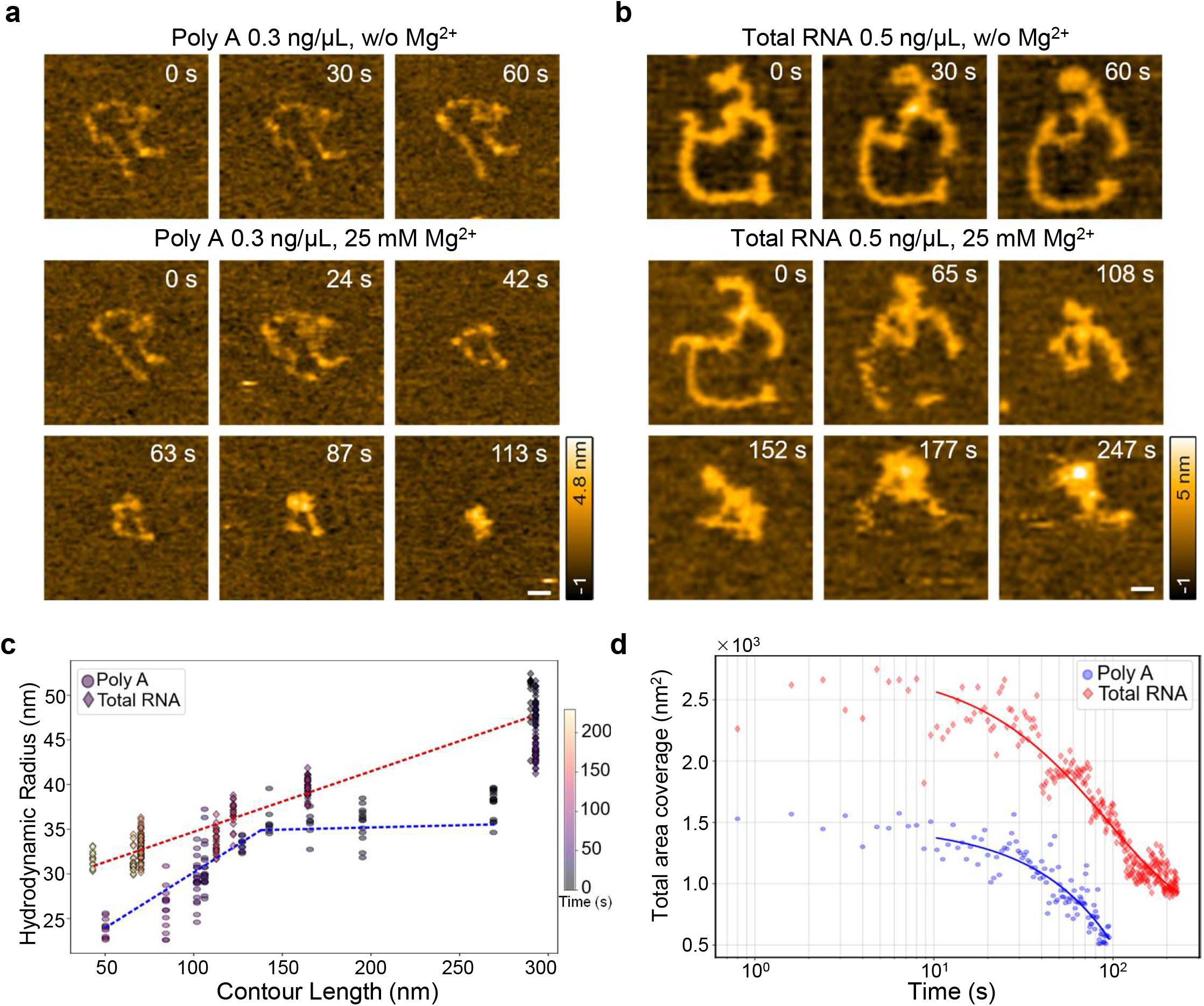
HS-AFM observation of single-molecule poly A and total RNA structural changes in Mg2+. a. Successive HS-AFM images showing the effects of Mg2+ on a single-stranded poly A. RNA molecule fold and formed globular shape in the presence of Mg2+. Experiments were conducted within a 350 nm x 350 nm scanning area with 120 x 120 pixels (Scale bar, 50 nm). b. Successive HS-AFM images showing real-time structural changes of individual total RNA molecules upon Mg2+ addition. The RNA molecules gradually folded and adopted globular conformations following Mg2+ addition. Experiments were conducted within a 350 nm × 350 nm scanning area at 120 × 120 pixels (Scale bar, 50 nm). c. Scaling of hydrodynamic radius with RNA contour length. Hydrodynamic radius (R) is estimated as the square root of the coverage area. R is plotted against contour length (L) for poly A and total RNA, with points colored by time. Dashed lines indicate linear regression fits (R² > 0.9) and are shown as trends. d. Time-dependent compaction and maturation of RNA condensates. Shrinkage of total area coverage is a direct measure of condensation via compaction of a single chain. Time evolution of area was analyzed to identify the formation and growth of the condensate.

To link single-molecule folding dynamics with emergent condensate properties, we next quantified the compaction and structural evolution of individual RNA molecules in real time. Hydrodynamic radius (*R*_H_), estimated from the projected area of individual RNA molecules, provides a quantitative measure of molecular compaction during condensation. R_H_ was plotted against contour length (L_C_) for both poly A and total RNA, colored according to the recording time (Figure 6c). Large R_H_ and L_C_ values correspond to early, extended conformations, while small R_H_ and L_C_ values indicate late-stage, compact structures. Discrete changes in L_C_ mark stepwise compaction along the polymer, whereas horizontal or nearly constant R_H_ at a given L_C_ (Plateau-like behavior) reflects intramolecular rearrangements without immediate size reduction. Although poly A and total RNA have comparable contour lengths (250–300 nm), their geometric evolutions differ. Poly A initially exhibits plateau-like behavior, with R_H_ remaining relatively constant over a range of L_C_ values, followed by rapid compaction into condensed droplets. This reflects uniform base-stacking interactions and isotropic molecular behavior. In contrast, total RNA shows a positive linear correlation between R_H_ and L_C_, indicating gradual, stepwise compaction with concurrent structural rearrangements, consistent with a heterogeneous charge distribution and structured regions guiding the molecule through multiple conformational basins. Shrinkage of total area coverage further quantifies single-chain condensation (Figure 6d). Here, the projected area of individual RNA molecules was tracked over time, providing a direct measure of molecular compaction. Early time points (∼0–10 s) show large areas corresponding to extended conformations. Poly A molecules compact rapidly, with nearly linear area reduction, consistent with smooth droplet-like condensation. In contrast, total RNA exhibits slower and exponential area decrease, reflecting stepwise folding and rearrangements within structurally heterogeneous sequences. Compaction begins for both RNA types around 10 s, but the maturation kinetics differ markedly: Poly A evolves with a rapid linear rate (∼0.1 mn^2^.s^-1^), while total RNA grows more slowly and exponentially (∼0.01 nm^2^.s^-1^) (Figure 6d), indicating distinct energy landscapes and folding pathways.

These analyses reveal that the uniform backbone and base-stacking interactions of poly A enable isotropic molecular interactions, promoting rapid and smooth compaction. In contrast, total RNA, with its heterogeneous sequence and structured regions, undergoes irregular, stepwise condensation, producing non-spherical assemblies with limited coalescence. Together with relaxation time measurements, these data provide a quantitative readout of material properties, distinguishing droplet-like, liquid behavior of poly A from aggregate-like, solid behavior of total RNA.

### Chelation-Induced Collapse of RNA Condensates Highlights the Structural Role of Mg^2+^

Divalent cations potentially function both in the formation of condensates by promoting RNA folding and intermolecular interactions, and in the maintenance of their structural integrity once assembled. To further investigate the role of Mg^2+^ in condensate stability, we tested the reversibility of RNA condensation by chelating Mg^2+^ with ethylenediaminetetraacetic acid (EDTA). Due to its high affinity for divalent cations, EDTA effectively sequesters Mg^2+^ in a 1:1 stoichiometry, forming highly stable complexes independent of the cationic charge^[36–38]^. To test the reversibility of RNA condensation, 5 ng/µL of each RNA was incubated in a 20 µL reaction containing 50 mM MgCl₂ and 10 mM Tris-HCl (pH 7.4) at room temperature for 15 minutes. A 2 µL aliquot was then subjected to HS-AFM imaging. Before adding EDTA, fine circular droplets of poly A (Figure 7a & Movie 6) and various-sized total RNA aggregates (Figure 7b & Movie 7) were observed in HS-AFM. While imaging, EDTA was added to the scanning chamber to chelate Mg^2+^ ions in real time. Within ten minutes of EDTA addition, poly A droplets exhibited signs of structural disintegration (Figure 7a & Movie 8). The area of poly A droplets decreased within the first 10 minutes (Figure7c), likely reflecting the release of RNA molecules from larger droplets, resulting in smaller droplets. Subsequently, these smaller droplets appeared to unfold into linear RNA between 10 and 20 minutes, causing an increase in area (Figure 7c). Height and circularity decreased continuously over this period (Figure 7d, e).

**Figure 7.**
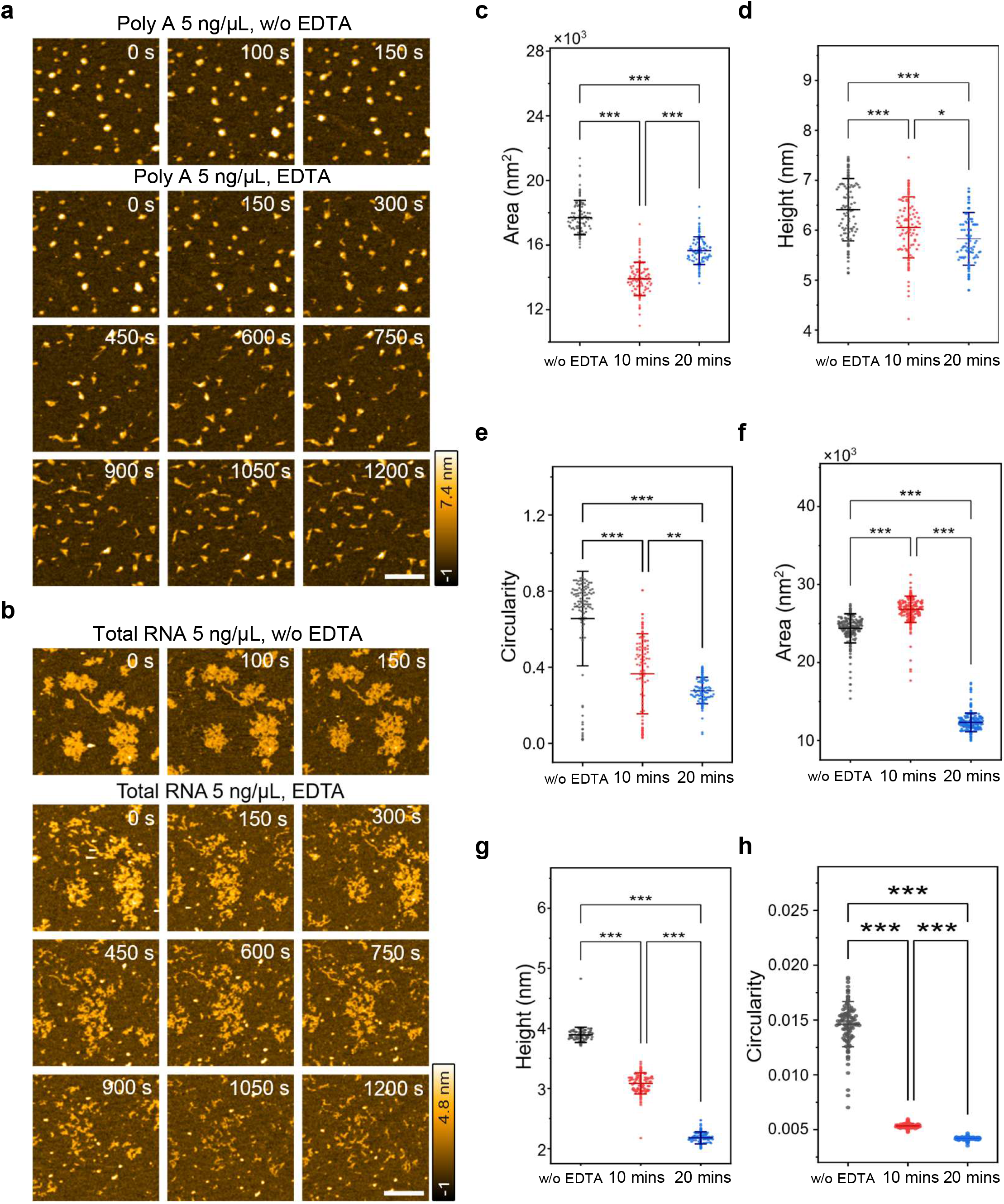
EDTA-induced structural relaxation of Mg2+ dependent RNA droplets and aggregates. a. Successive HS-AFM images showing the effect of EDTA on poly A droplets. Within 10 mins of EDTA addition, poly A droplets begin to lose their integrity and shape, indicating partial structural relaxation. Experiments were conducted within a 500 nm x 500 nm scanning area with 120 x 120 pixels (Scale bar, 100 nm). b. HS-AFM observation of total RNA aggregates in the presence of Mg2+. The addition of EDTA gradually reduces the integrity and size of condensations. Experiments were conducted within a 500 nm x 500 nm scanning area with 120 x 120 pixels (Scale bar 100 nm). c. d. e. Scatter dot plots showing the change of area, height and circularity of poly A droplets over time following EDTA addition. One-way ANOVA test confirmed significant difference between groups (n=100, *; p<0.05. **; p<0.01, ***; p < 0.001). f. g. h. Scatter dot plots showing the changes in area, height and circularity of total RNA condensation after EDTA addition. A one-way ANOVA test was performed to compare the means at different incubation times. (n=100, *; p<0.05. **; p<0.01, ***; p < 0.001).

In contrast to poly A RNA, the area of total RNA increased during the first 10 minutes, presumably due to the unfolding of aggregates, and then decreased by 20 minutes as unfolded RNA molecules were released (Figure 7b, f, Movie 9). This behavior supports the reversibility of the assembly process, in which intramolecular interactions initially cause size expansion, followed by structural rearrangements that lead aggregates to adopt tighter conformations and reduce their size (Figure 3a). The slower disassembly dynamics of total RNA compared to poly A RNA, which rapidly adjusts droplet cluster size via diffusion, further support the occurrence of ongoing structural transitions in total RNA after intermolecular interactions. The height of total RNA decreased upon EDTA addition (Figure 7g), while the circularity of total RNA aggregates decreased aggressively during the first 10 minutes and then slightly decreased from 10 to 20 minutes, as unfolding of RNA into linear forms would be expected to reduce circularity (Figure 7h). This may reflect the loss of loosely associated or surface-bound RNA molecules during diffusion, leaving behind a population of smaller RNAs. Together, these results support distinct condensation behaviors between poly A and total RNA, suggesting that stabilizing interactions such as π–π stacking and other multivalent contacts, independent of RNA–RNA base pairing, enable rapid, RNA-dependent droplet formation.

## Discussion

RNAs act as scaffolds that facilitate LLPS, primarily through the formation of RNA structures and RNA–RNA interactions, which play central roles in RNA-driven phase separation^[31,39,40]^. In this study, we introduce a powerful approach to directly visualize dynamic transitions of RNA-driven LLPS at the single-molecule level using HS-AFM. This approach enables real-time monitoring of RNA molecular conformations, intermolecular interactions, and cluster merging dynamics, providing the first single-molecule visualization of RNA condensation dynamics in real time. A major challenge in studying RNA-driven LLPS with HS-AFM lies in balancing target molecule concentrations. Because HS-AFM relies on scanning-based detection, it requires lower RNA concentrations, whereas LLPS typically occurs at higher concentrations. By carefully optimizing Mg^2+^ and RNA levels, we established experimental conditions that enabled direct visualization of RNA transitions from linear molecules to self-folded structures, cluster formation, and larger condensates while maintaining single-molecule resolution. Importantly, the condensation observed by HS-AFM was consistent with DLS measurements performed in bulk solution, demonstrating that the observed size evolution and assembly dynamics are not artifacts of surface confinement. This agreement between surface-based imaging and solution-phase measurements indicates that surface effects are minimal under our conditions and supports the accuracy of our real-time observations. Taken together, this approach provides a powerful platform to dissect the mechanistic steps of RNA condensation in real time.

Previous studies have shown that high concentrations of Mg^2+^ or Ca^2+^ can induce condensation of homopolymeric RNAs such as poly U ^[40]^, and that RNA-driven condensates can recapitulate key features of membraneless organelles in cells^[14]^. We demonstrated that RNA condensation occurs through intermolecular RNA–RNA interactions in the presence of divalent cations (Mg^2+^), independent of crowding agents, consistent with previous studies^[41,42]^. Building on these findings, we extended our analysis to both homopolymeric poly A and total RNA pools (synthetic and natural RNA), as also demonstrated in other approaches^[31]^, to assess whether RNA condensation is a generalizable and biologically relevant phenomenon. Interestingly, poly A RNA can rapidly and efficiently form droplets at room temperature without heat, despite the absence of canonical base-pairing. This highlights a critical point: intrinsic RNA properties that modulate condensation are not limited to base-pairing interactions but also involve other molecular interactions. In contrast, total RNA initially forms aggregates rather than droplet-like condensates without heat, suggesting that sequence composition is essential.

Our findings support a hierarchical model of RNA condensation in which intramolecular folding precedes and facilitates intermolecular RNA–RNA interactions. High-resolution HS-AFM imaging directly visualizes this progression, with single-stranded RNA first forming secondary structures via cis interactions in the presence of Mg²⁺ before engaging in higher-order assembly. Overall, poly A and total RNA undergo condensation through fundamentally distinct folding and merging dynamics (Figure 8, Table 1). These differences likely arise from intrinsic disparities in sequence composition and structural complexity. As a homopolymer, poly A can engage in relatively uniform stacking interactions that promote cooperative and isotropic molecular collapse. In contrast, total RNA contains heterogeneous sequences and performed secondary structures, introducing structural anisotropy and competing interaction modes that constrain molecular reorganization. Such molecular-level properties propagate to mesoscale assembly behavior. Poly A assemblies readily reorganize upon contact, enabling efficient coalescence into smooth, spherical droplets whose growth is dominated by fusion. By contrast, total RNA clusters appear to undergo continued internal rearrangement after association, reflecting competition between intra- and intermolecular interactions. This hierarchical reorganization likely underlies the non-monotonic size evolution observed during cluster maturation and promotes the formation of structurally constrained assemblies.

**Figure 8.**
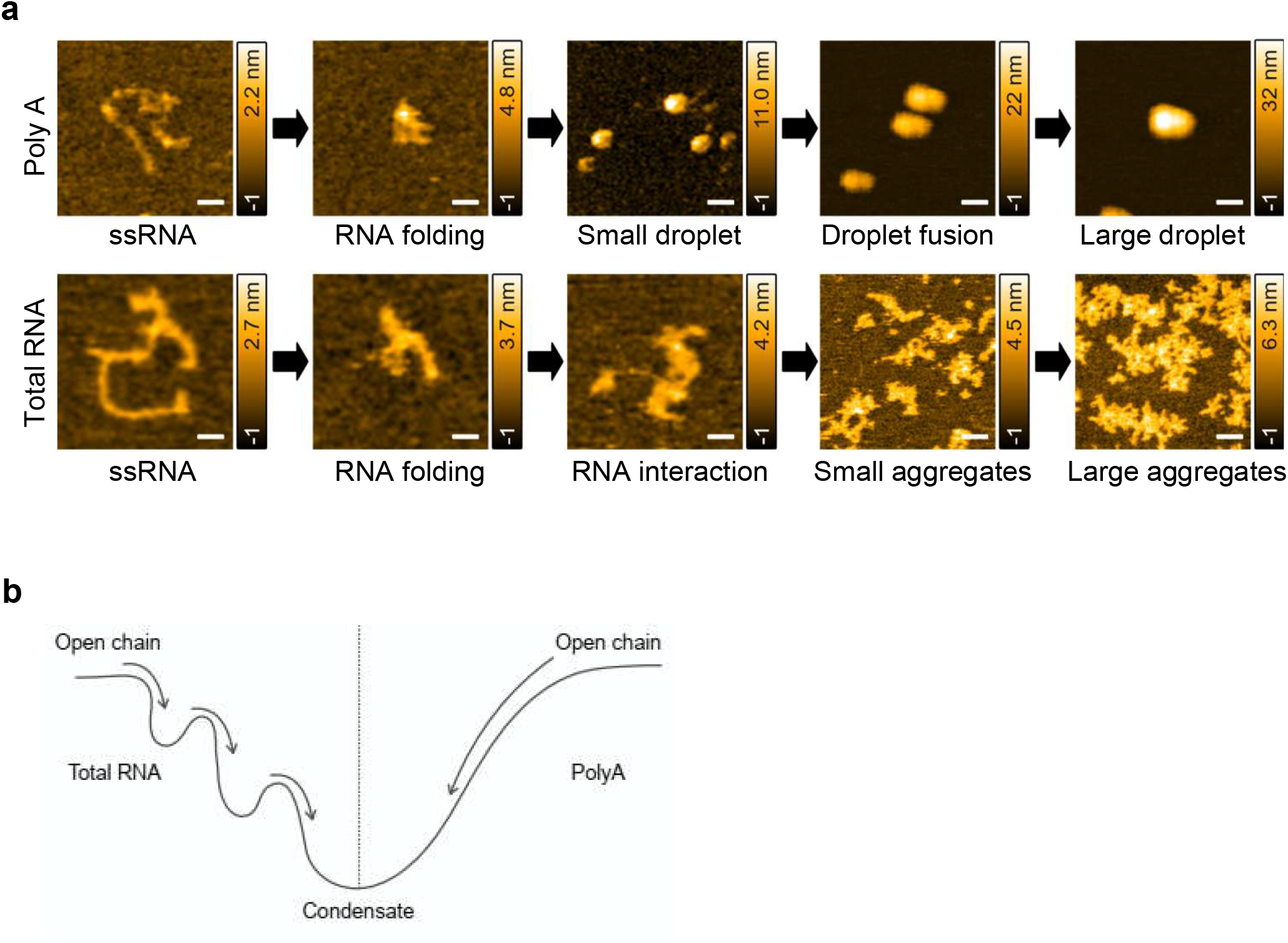
Overview of RNA condensation maturation. a. Stepwise condensation poly A (droplet-like) and total RNA (aggregation-like) during Mg²⁺ induced condensate formation (Scale bar, 50 nm). b. Schematic of condensation maturation in poly A and total RNA. Poly A exhibits a two-state energy landscape with open and condensed states. After a brief rearrangement phase with minimal change in hydrodynamic radius (R), poly A rapidly collapses into the condensed state. In contrast, total RNA follows a multi-step, corrugated energy landscape with gradual maturation, reflecting the difference between droplet-like and aggregation-like condensation.

**Table 1.** Physiological properties of poly A and total RNA condensates.

| **Features** | **Poly A** | **Total RNA** |
| --- | --- | --- |
| Morphology | Droplet like | Solid/gel like aggregate |
| Shape | Circular | Irregular |
| PDI | 0.33 (<0.5) | 0.69 (>0.5) |
| Height | ~ 8 nm or more | Below 8 nm |
| Circularity | 0.5 > | 0.5 < |
| Young's modulus | 3–10 MPa | 10–40 MPa |
| Indentation depth | High | Low |
| Relaxation time ($\tau$) | 1.63 s | 8.32 s |

Mechanical measurements further support the distinct material properties of these RNA condensates. Greater indentation depth in poly A condensates indicates higher deformability, consistent with droplet-like behavior. In contrast, total RNA assemblies exhibit higher Young’s modulus, reflecting greater resistance to deformation and a more rigid, aggregate-like state. These findings suggest that RNA sequence complexity and folding heterogeneity shape the condensation energy landscape, biasing assemblies toward either dynamic liquid-like states or more rigid solid-like states. In this framework, RNA structure is not merely permissive for phase separation but acts as a key determinant of the material properties of RNA-driven condensates. In future studies, this platform can be extended to systematically dissect how defined RNA sequences, biologically relevant ions (e.g., K⁺, Na⁺, Ca^2+^), and protein factors, both individually and in combination, influence RNA folding landscape and interaction networks. Such analyses will clarify how cis-encoded structural features and trans-acting modulators collectively determine the emergent material state of RNA assemblies. Integrating these parameters will refine biophysical models of intracellular phase transitions and help bridge controlled in vitro observations with the complexity of living cells. These insights are particularly relevant for RNA-rich membraneless organelles, such as SG, P-bodies, and nucleoli, which respond dynamically to cellular stress, signaling pathways, and gene expression programs. More broadly, defining how sequence and structure dependent RNA condensation governs material properties has important biomedical implications, as dysregulated RNA aggregation and aberrant phase transitions are increasingly linked to neurodegeneration, cancer, and other pathologies. Altogether, this strategy opens new avenues connecting RNA sequence and folding complexity to condensate dynamics, material states, and cellular function.

## Materials and Methods

### Isolation of RNA from HEK293T cells

Total RNA was extracted from human HEK293T cells using the NucleoSpin RNA Plus kit (Macherey-Nagel, Germany) according to the manufacturer’s protocol. RNA quality was checked using agarose gel electrophoresis. Poly A RNA was purchased from Roche (catalog number 10108626001).

### In vitro condensation experiments and Bright-Field (BF) Imaging

For the total RNA, purified RNA samples were resuspended in nuclease-free water to desired final concentration. RNA was then mixed with 10 mM Tris-HCl (pH 7.4) and varying concentrations of MgCl_2_ as specified. For the experiments shown in Figure S1, the mixture contained 150 ng/µL total RNA, 10% PEG (MW3350),10 mM MgCl_2_, and 10 mM Tris-HCl (pH 7.4). A total volume of 20 µL of each mixture was prepared in a tube and subjected to the following heat treatment: samples were heated at 90°C for 3 minutes, then transferred to 65°C for 1 minute, and finally incubated at 37°C until imaging. For the poly A RNA, the same experimental procedure was followed as for total RNA. For bright-field imaging, 10 µL of each sample was placed on a coverslip and observed using Keyence BZ-X810 microscope equipped with a 60× objective lens. Images were acquired using the BZ-X800 image analyzer and processed with the ImageJ Fiji software package ^[43]^.

### Turbidity Measurement

To measure the turbidity 100 µL solution of both RNAs were prepared in varied Mg^2+^ concentrations and measured in quartz cuvette. The solutions were mixed and injected carefully into cuvette. The absorbance of the solution was measured 300-400 nm for 1 h on Jasco V-570 spectrophotometer at 25°C with shaking. The experiments were conducted for three independent samples and mean ± SD values were calculated.

### Dynamic light scattering measurements

The dynamic light scattering experiment was carried out on a Malvern Zetasizer nano series instrument. RNA solutions of 100 µL with defined concentration of Mg^2+^ were measured in clear quartz cuvette. The intensity of scattered light was recorded at a scattering angle of 90° at 25°C for 1 h collecting 10 acquisitions (8s each). Each experiment was repeated 3 times. Zetasizer software 7.12 was used to analyze the particle size.

### High-speed AFM imaging

A laboratory-built high-speed atomic force microscope^[44]^ was used as described previously^[45]^. We used small cantilevers (BLAC10DS-A2, Olympus, Japan) with a nominal spring constant of ∼0.1 N/m, resonant frequency of ∼0.5 MHz, and a quality factor of ∼1.5 in liquid. An amorphous carbon tip was fabricated on the original AFM tip by electron beam deposition. The length of the additional AFM tip was ∼500 nm, and the tip was further sharpened using a radio-frequency plasma etcher (Tergeo, PIE Scientific LLC, USA) under an argon gas atmosphere (Direct mode, 10 sccm, and 20 W for 1.5 min). Regarding the sample deposition, a glass sample stage (diameter, 2 mm; height, 2 mm) with a thin mica disc (1.5 mm in diameter and ∼0.05 mm in thickness) glued to the top by epoxy was attached onto the top of a Z-scanner by a drop of nail polish.

As a substrate, 3-aminopropyltriethoxysilane (APTES) treated mica was used only for single molecule imaging (Figure 6a,b) as described previously ^[46]^. APTES was diluted 10^6^ times with MilliQ-water, and 2 µL of the fresh solution was applied to the mica surface for 3 minutes. After rinsing the surface five times with MilliQ-water of 20 µL, the surface was further rinsed with the imaging buffer (10 mM Tris-HCl, pH 7.5, 20 mM NaCl, 60 mM KCl) of 20 µL, during which the surface was not allowed to dry. Then, 2 µL of RNA sample diluted with the imaging buffer was applied to the surface for 3 minutes. After rinsing the surface with the imaging buffer of 20 µL, the surface was immersed in the imaging chamber containing the imaging buffer of 60 µL and then imaged with HS-AFM in the tapping mode. The free oscillation amplitude of the cantilever was 2-3 nm, and the set-point amplitude was set to 90% of the free amplitude. MgCl_2_ was added to the imaging chamber to obtain the desired final concentrations. For Mg^2+^ chelation experiments, RNA condensates were first performed in a tube containing 5 ng/µL RNAs and 25 mM Mg^2+^. Subsequently, 2 µL of the RNA solution was introduced into the imaging chamber containing scanning buffer, resulting in dilution of Mg^2+^ to approximately 0.7 mM. To chelate Mg^2+^, EDTA was then added to a final concentration of 10 mM, sufficient to remove all remaining free Mg^2+^ ions under these conditions.

All HS-AFM experiments were performed at 24-26 °C, and data were collected using laboratory-developed software UMEX. Image contrast was normalized as needed for visualization. The height, circularity, and area of objects in HS-AFM images were measured and analyzed using UMEX Viewer. Linear RNA length was determined using the line analysis function in UMEX Viewer, and the longest dimension of globular RNAs was measured as their length after folding.

### Quantification of coalescence and shape relaxation kinetics

Relaxation kinetics following coalescence were quantified using time-resolved circularity measurements. Coalescence events were identified from HS-AFM time-lapse imaging, and the onset of merging between two assemblies was defined as t = 0.

Circularity was calculated for each condensate at successive time points using measurements derived from the radius of gyration. Circularity recovery was monitored from t = 0 to 30 s post-coalescence, as merged assemblies reached morphological stabilization within this time window. The relaxation time constant (τ) was determined by fitting the circularity recovery curve to a single-exponential model:

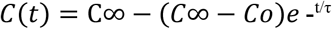

where:

- *C*(*t*)is the circularity at time *t*,
- *C*_0_is the circularity at the onset of coalescence (t = 0),
- *C*_∞_is the plateau circularity after stabilization, and
- *ô* represents the characteristic relaxation time constant.

For each sample, τ values were calculated from multiple independent coalescence events and averaged to obtain the mean relaxation time.

### Kinetic Analysis of Single-Molecule RNA Folding

Hydrodynamic radius estimation: The hydrodynamic radius (R_H_) of individual RNA molecules was estimated as the square root of the projected area measured from HS-AFM images. For single chains of charged monomers, this approximation is valid based on the formal electrostatics– hydrodynamics analogy.

Contour length tracking and R_H_–L_C_ analysis: The contour length (L_C_) of each molecule was determined from traced RNA backbones. R_H_ was plotted against L_C_ for both poly A and total RNA, with points colored according to the time of recording. Linear regression was applied to the R_H_– L_C_ data (goodness-of-fit >90%) to quantify trends. Vertical clustering of data (change in R_H_ at nearly constant L_C_) was used to identify potential intramolecular structural rearrangements, whereas discrete decreases in L_C_ were used to indicate stepwise chain compaction.

Area coverage monitoring: The projected area of individual RNA molecules was tracked over time to quantify single-chain condensation. The shrinkage of total area coverage was used as a direct metric of compaction, allowing identification of initial folding, intermediate rearrangements, and subsequent condensate growth.

Time resolution and growth rate measurement: Compaction dynamics were monitored with a temporal resolution sufficient to capture early events (∼10 s post Mg²⁺ addition). Growth rates of the area coverage were estimated for both poly A and total RNA by evaluating the time-dependent change in projected area, enabling comparison of linear versus exponential compaction behaviors.

### 3D Force-Mapping Nanomechanical Measurements

The 3D force-mapping measurements were performed using the laboratory-developed software UMEX Sample-Scan HS-AFM, employing the same cantilever used for topographic imaging. With cantilever excitation turned off, the probe was approached toward the surface in the z direction while monitoring the deflection, and it was retracted when the deflection reached a value corresponding to ∼0.8 nN. The approach and retraction velocities were both set to 2.9 µm/s. The scan area in the XY plane was 70 × 70 nm^2^ (40 × 40 pixels) for poly A droplets and 150 × 150 nm^2^ (40 × 40 pixels) for total RNA aggregates. The Z-range was 20–30 nm (≈110 pixels) in both cases. Under these conditions, acquisition of a single 3D dataset required approximately 17 s. Reconstruction of topographic images and analysis of Young’s modulus and indentation depth from the 3D datasets were performed using the UMEX 3D Force Map Viewer.

Young’s modulus was calculated by fitting the force curve obtained at each pixel to the Hertz contact model assuming sphere–sphere contact^[47,48]^, as follows:

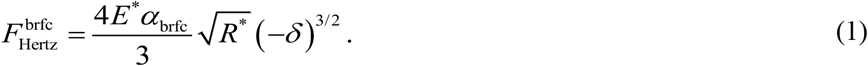

Here, *R*^∗^ denotes the effective radius of curvature, given by

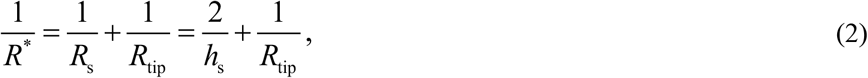

where *R*_tip_ represent the curvature radius of the probe tip and the condensate’s radius, which was assumed to be 10 nm. *R*_s_ and *h*_s_ represent the condensates radius and height, respectively, and the *h*_s_ was estimated from the surface topography acquired immediately prior to the 3D force-mapping measurement.

The parameter *E*^∗^ represents the reduced Young’s modulus. When the Young’s modulus of the probe is sufficiently larger than that of the sample, *E*^∗^ can be approximated by

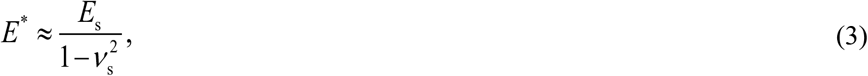

Where, E_s_ denotes the Young’s modulus of the sample, and ν_s_ denotes the Poisson’s ratio of the sample, which was assumed to be 0.5, corresponding to typical values for liquids and gels.

When the condensates thickness becomes small, the underlying substrate causes the apparent condensates modulus to be overestimated due to the bottom effect. Consequently, a correction was applied using the following equation^[47,48]^:

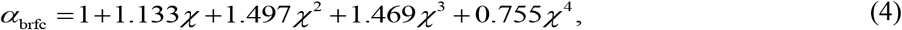

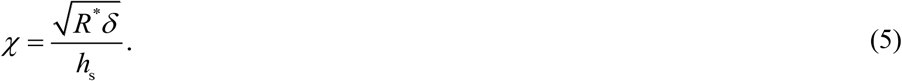

It should be noted that the sphere–sphere contact model described in Eq. (1) was applied uniformly to all pixels; therefore, modulus maps shown in (Figure 5 e,f) obtained on nominally flat substrate regions are not quantitatively accurate.

In the statistical analysis shown in (Figure 5 i,j) the Young’s modulus and indentation depth values for each dataset were averaged on logarithmic and linear scales, respectively. Both quantities exhibited weak correlations with *h*_s_, which may be attributed to the fact that Eq. (4) was derived under the assumption of a flat film and therefore may not fully compensate for substrate effects in spherical structures.

To determine whether statistically significant differences existed between the two different RNA condensates after accounting for *h*_s_ dependence, an analysis of covariance (ANCOVA) was performed. The *h*_s_ was treated as a covariate, and a dummy variable *D* (droplets = 0, aggregates = 1) was introduced to represent the condensate type. The Young’s modulus data were well described by

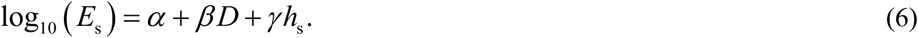

The regression assumed equal slopes and equal variance for both samples. The estimated coefficients were *α* = 1.411, *β* = 0.4126, and *γ* = −0.02781. ANCOVA indicated a significant effect of sample structure (*F* = 89.31, *p* = 4.857×10^−13^).

Similarly, for indentation depth, the data were fitted using

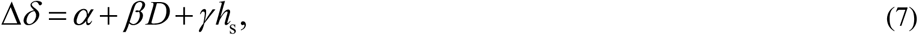

with estimated coefficients *α* = 1.087, *β* = −0.5676, and *γ* = 0.2888. The ANCOVA likewise showed a significant structural effect (*F*= 16.91, *p* = 1.343×10^−4^).

### Quantification and statistical analysis

All statistical analysis was performed in Origin Pro 2025b software. A significant cutoff value p < 0.05 was used for all analyses unless otherwise noted. The normality of distribution was tested by the Shapiro-Wilk test. To observe the mean difference among different droplet groups, a One-way ANOVA test was performed where the Tukey test was used for mean comparison.

## Data availability

The data and analysis code generated in this study are available from the corresponding authors upon reasonable request.

## Acknowledgments

We thank members of the Sato laboratories for their valuable discussions. We thank Prof. Toshio Ando, Ms. Kayo Nakatani and Ms. Risa Omura for the technical support of HS-AFM experiments.

## Author contributions

Conceptualization: HS

Methodology: SNM, NS, NK, HS

Investigation: SNM

Formal analysis: SNM

Data analysis & Visualization: SNM, NK, HS, KU, TD

Funding acquisition: SNM, NK, HS

Project administration: SNM, NK, HS

Supervision: NK, HS

Writing – original draft: SNM

Writing – review & editing: NK, KU, NS, HS

## References

[1] B. Wang, L. Zhang, T. Dai, Z. Qin, H. Lu, L. Zhang, F. Zhou, Signal Transduct. Target. Ther. 2021, 6, DOI 10.1038/s41392-021-00678-1.

[2] Y. S. Mao, B. Zhang, D. L. Spector, Trends in Genetics 2011, 27, 295.

[3] Z. Xu, W. Wang, Y. Cao, B. Xue, Supramolecular Materials 2023, 2, DOI 10.1016/j.supmat.2023.100049.

[4] L. Tang, Nat. Methods 2019, 16, 18.

[5] S. F. Banani, H. O. Lee, A. A. Hyman, M. K. Rosen, Nat. Rev. Mol. Cell Biol. 2017, 18, 285.

[6] Y. Shin, C. P. Brangwynne, Science (1979). 2017, 357, DOI 10.1126/science.aaf4382.

[7] J. Wang, J. M. Choi, A. S. Holehouse, H. O. Lee, X. Zhang, M. Jahnel, S. Maharana, R. Lemaitre, A. Pozniakovsky, D. Drechsel, I. Poser, R. V. Pappu, S. Alberti, A. A. Hyman, Cell 2018, 174, 688.

[8] L. P. Bergeron-Sandoval, N. Safaee, S. W. Michnick, Cell 2016, 165, 1067.

[9] Y. L. P. Ow, D. R. Green, Z. Hao, T. W. Mak, Nat. Rev. Mol. Cell Biol. 2008, 9, 532.

[10] X. Tong, R. Tang, J. Xu, W. Wang, Y. Zhao, X. Yu, S. Shi, Signal Transduct. Target. Ther. 2022, 7, DOI 10.1038/s41392-022-01076-x.

[11] P. Yang, C. Mathieu, R. M. Kolaitis, P. Zhang, J. Messing, U. Yurtsever, Z. Yang, J. Wu, Y. Li, Q. Pan, J. Yu, E. W. Martin, T. Mittag, H. J. Kim, J. P. Taylor, Cell 2020, 181, 325.

[12] J.-L. Liu, J. G. Gall, U Bodies Are Cytoplasmic Structures That Contain Uridine-Rich Small Nuclear Ribonucleoproteins and Associate with P Bodies, 2007.

[13] J. R. Buchan, R. Parker, J. Ross, Eukaryotic Stress Granules: The Ins and Out of Translation, n.d.

[14] N. Kedersha, M. D. Panas, C. A. Achorn, S. Lyons, S. Tisdale, T. Hickman, M. Thomas, J. Lieberman, G. M. McInerney, P. Ivanov, P. Anderson, Journal of Cell Biology 2016, 212, 845.

[15] B. Van Treeck, D. S. W. Protter, T. Matheny, A. Khong, C. D. Link, R. Parker, Proc. Natl. Acad. Sci. U. S. A. 2018, 115, 2734.

[16] M. M. Fay, P. J. Anderson, J. Mol. Biol. 2018, 430, 4685.

[17] Q. Guo, X. Shi, X. Wang, Noncoding RNA Res. 2021, 6, 92.

[18] Y. Ma, H. Li, Z. Gong, S. Yang, P. Wang, C. Tang, J. Am. Chem. Soc. 2022, 144, 4716.

[19] I. S. Tolokh, S. A. Pabit, A. M. Katz, Y. Chen, A. Drozdetski, N. Baker, L. Pollack, A. V. Onufriev, Nucleic Acids Res. 2014, 42, 10823.

[20] A. Jain, R. D. Vale, Nature 2017, 546, 243.

[21] M. M. Fay, P. J. Anderson, P. Ivanov, Cell Rep. 2017, 21, 3573.

[22] W. Ma, G. Zheng, W. Xie, C. Mayr, Elife 2021, 10, DOI 10.7554/eLife.64252.

[23] B. Drobot, J. M. Iglesias-Artola, K. Le Vay, V. Mayr, M. Kar, M. Kreysing, H. Mutschler, T. Y. D. Tang, Nat. Commun. 2018, 9, DOI 10.1038/s41467-018-06072-w.

[24] C. Ehresmann, F. Baudin, M. Mougel, P. Romby, J.-P. Ebel, B. Ehresmann, Probing the Structure of RNAs in Solution, n.d.

[25] D. J. Müller, A. C. Dumitru, C. Lo Giudice, H. E. Gaub, P. Hinterdorfer, G. Hummer, J. J. De Yoreo, Y. F. Dufrêne, D. Alsteens, Chem. Rev. 2021, 121, 11701.

[26] E. P. Holmes, M. C. Gamill, J. I. Provan, L. Wiggins, R. Rusková, S. Whittle, T. E. Catley, K. H. S. Main, N. Shephard, H. E. Bryant, N. S. Gilhooly, A. Gambus, D. Račko, S. D. Colloms, A. L. B. Pyne, Nature Communications 2025, 16, DOI 10.1038/s41467-025-60559-x.

[27] G. R. Heath, S. Scheuring, Curr. Opin. Struct. Biol. 2019, 57, 93.

[28] T. Ando, S. Fukuda, K. X. Ngo, H. Flechsig, 2025, 158, 12.

[29] Y. L. Lyubchenko, L. S. Shlyakhtenko, T. Ando, Methods 2011, 54, 274.

[30] O. Annunziata, N. Asherie, A. Lomakin, J. Pande, O. Ogun, G. B. Benedek, Effect of Polyethylene Glycol on the Liquid-Liquid Phase Transition in Aqueous Protein Solutions, n.d.

[31] E. N. H. Phan, C. H. Mak, 2025, DOI 10.1261/rna.

[32] V. K. Misra, D. E. Draper, A Thermodynamic Framework for Mg 2 Binding to RNA A Conceptual Framework for Mg 2, n.d.

[33] J. Schauss, A. Kundu, B. P. Fingerhut, T. Elsaesser, Journal of Physical Chemistry B 2021, 125, 740.

[34] G. Feldmann, Z. E. Fabrim, G. L. Hennig, in J. Mater. Sci., 2008, pp. 614–620.

[35] A. Ghosh, D. Kota, H. X. Zhou, Nat. Commun. 2021, 12, DOI 10.1038/s41467-021-26274-z.

[36] N. A. Yewdall, A. A. M. André, T. Lu, E. Spruijt, Curr. Opin. Colloid Interface Sci. 2021, 52, DOI 10.1016/j.cocis.2020.101416.

[37] P. L. Onuchic, A. N. Milin, I. Alshareedah, A. A. Deniz, P. R. Banerjee, Sci. Rep. 2019, 9, DOI 10.1038/s41598-019-48457-x.

[38] R. Yamagami, J. P. Sieg, P. C. Bevilacqua, Biochemistry 2021, 60, 2374.

[39] J. K. A. Tom, P. L. Onuchic, A. A. Deniz, Journal of Physical Chemistry B 2022, 126, 9715.

[40] I. R. E. A. Trussina, A. Hartmann, C. Desroches Altamirano, J. Natarajan, C. M. Fischer, M. Aleksejczuk, H. Ausserwöger, T. P. J. Knowles, M. Schlierf, T. M. Franzmann, S. Alberti, Mol. Cell 2025, 85, 585.

[41] D. M. Parker, D. Tauber, R. Parker, Mol. Cell 2025, 85, 571.

[42] R. R. Poudyal, J. P. Sieg, B. Portz, C. D. Keating, P. C. Bevilacqua, 2021, DOI 10.1261/rna.

[43] J. Schindelin, I. Arganda-Carreras, E. Frise, V. Kaynig, M. Longair, T. Pietzsch, S. Preibisch, C. Rueden, S. Saalfeld, B. Schmid, J. Y. Tinevez, D. J. White, V. Hartenstein, K. Eliceiri, P. Tomancak, A. Cardona, Nat. Methods 2012, 9, 676.

[44] D. Kilburn, R. Behrouzi, H. T. Lee, K. Sarkar, R. M. Briber, S. A. Woodson, Nucleic Acids Res. 2016, 44, 9452.

[45] T. Uchihashi, N. Kodera, T. Ando, Nat. Protoc. 2012, 7, 1193.

[46] N. Kodera, H. Abe, P. D. N. Nguyen, S. Ono, Journal of Biological Chemistry 2021, 296, DOI 10.1016/j.jbc.2021.100649.

[47] V. G. Gisbert, R. Garcia, ACS Nano 2021, 15, 20574.

[48] P. D. Garcia, R. Garcia, Biophys. J. 2018, 114, 2923.

