## Supplementary figures for "HS-AFM Reveals Hierarchical RNA Folding Driving Condensate Assembly and Material States"

**Figure S1**

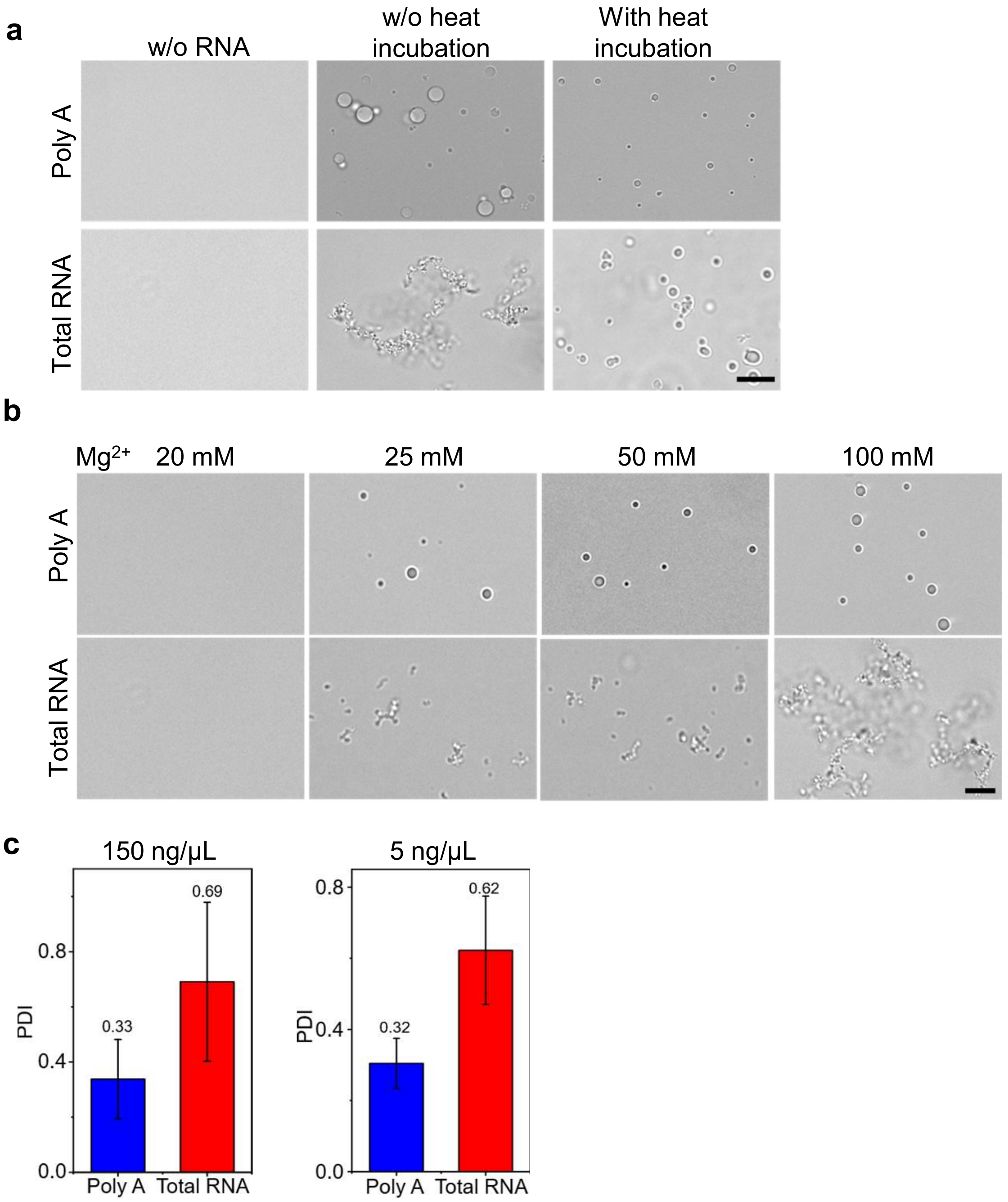


**Figure S1. Droplet formation of poly A and total RNA at different physiological conditions.**

a. Condensation of poly A and total RNA (150 ng/µL) in 10% PEG, 10 mM $MgCl_2$, 10 mM Tris-HCl (pH 7.4), with or without heat. Poly A RNA formed spherical droplets under both conditions, while total RNA formed aggregates without heat and droplet-like structures only after heat treatment (Scale bar 10 µm).

b. Both poly A (150 ng/µL) and total RNA (150 ng/µL) formed condensates at 20, 25, 50, 100 mM $Mg^{2+}$, where number and size increasing at higher $Mg^{2+}$ concentrations (Scale bar 10 µm).

c. Bar graph representing polydispersity index (PDI) of poly A droplets and total RNA aggregates at 150 ng//µL (Left panel) and 5 ng//µL (Right panel). Poly A droplets have lower PDI compared to total RNA aggregates in both conditions indicating poly A forms monodisperse droplets.

**Figure S2**

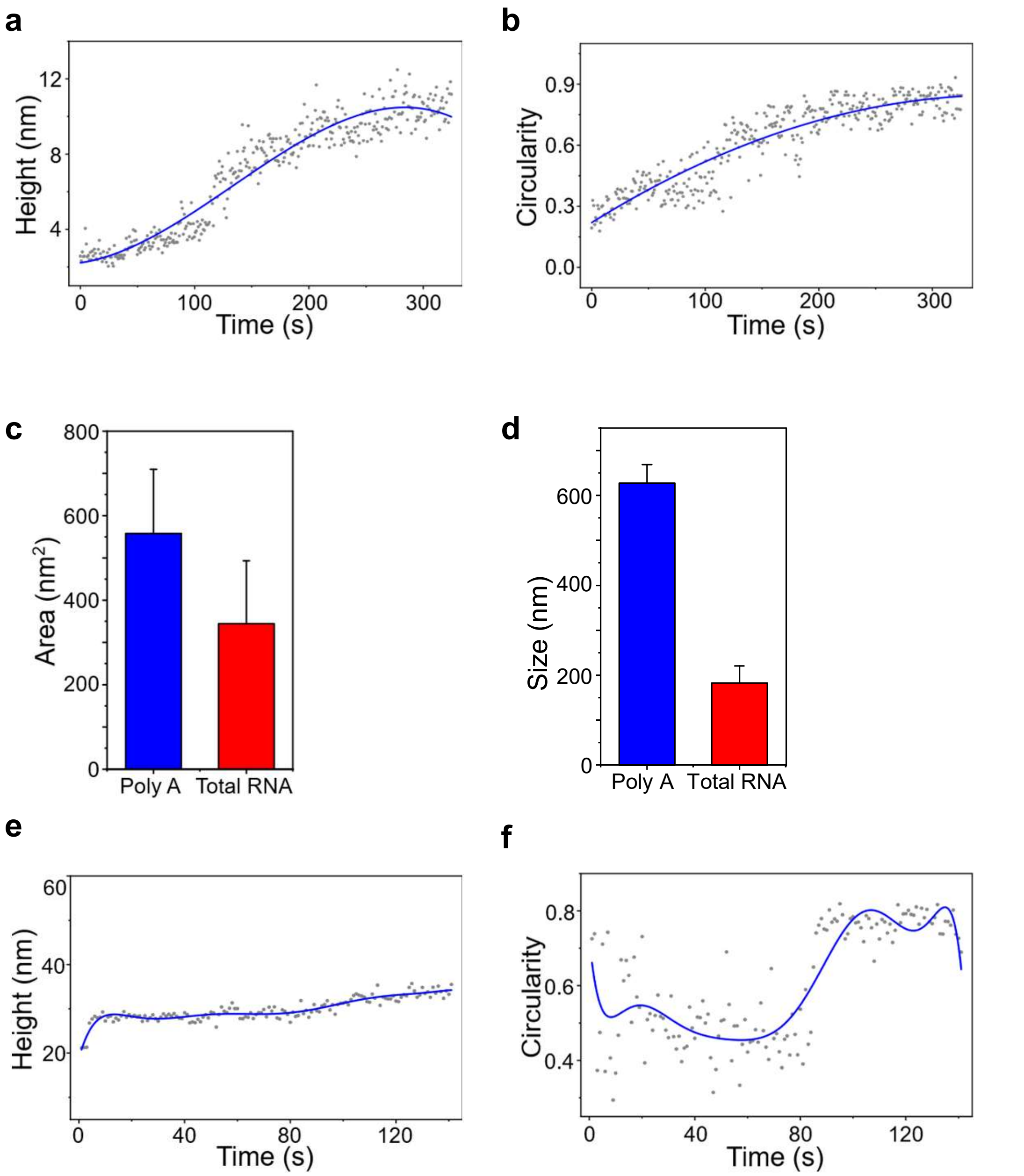


**Figure S2. Physiological properties of poly A droplets.**

**a.b.** Scatter plots showing changes in height, and circularity of droplets formed by poly A RNA in condition (2a). Height and circularity increased over time following $Mg^{2+}$ addition. The solid line represents a polynomial fit to the experiment data, illustrating the increasing trend of data during droplet formation.

**c.** Bar graph showing average area of condensates after 5 mins of $Mg^{2+}$ addition at HS-AFM in experiment condition (2a and 3a). Data was plotted mean ± SD of 20 samples of both poly A droplets and total RNA aggregates.

**d.** Bar graph showing average size (diameter) of condensates after 5 mins of $Mg^{2+}$ addition at DLS in similar experiment condition (2a and 3a). Data was plotted mean ± SD of three individual experiments (n=3) of both poly A droplets and total RNA aggregates.

**e.f.** Scatter plot showing changes in height and circularity during droplet maturation in condition (2c). Both height and circularity increase with time as droplets merge to form large droplets. The solid line represents a polynomial fit to the experiment data, illustrating the increasing trend of data during droplet maturation.

### Figure S3

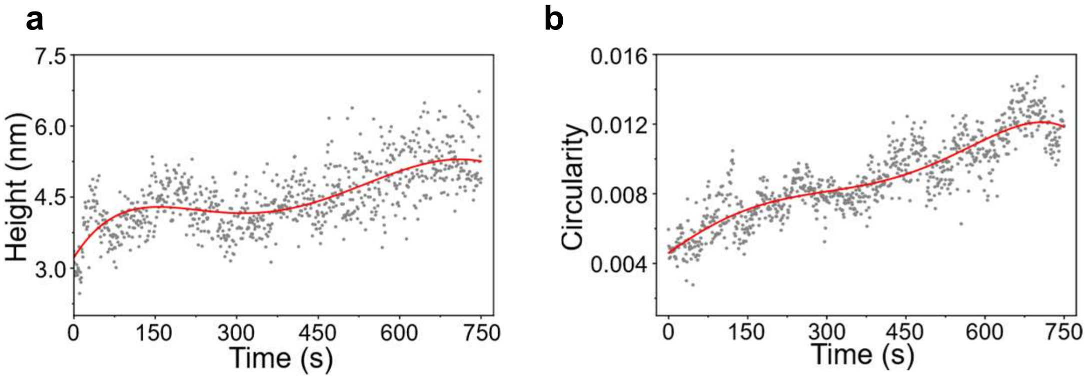


**Figure S3. Physiological properties of total RNA aggregates.**
**a.b.** Scatter plots showing changes in height, and circularity of total RNA during aggregate formation under the conditions in (3a). Height and circularity increased slightly over time. The solid line represents a polynomial fit to the experiment data, illustrating the increasing trend of data during aggregates formation.

**Figure S4**

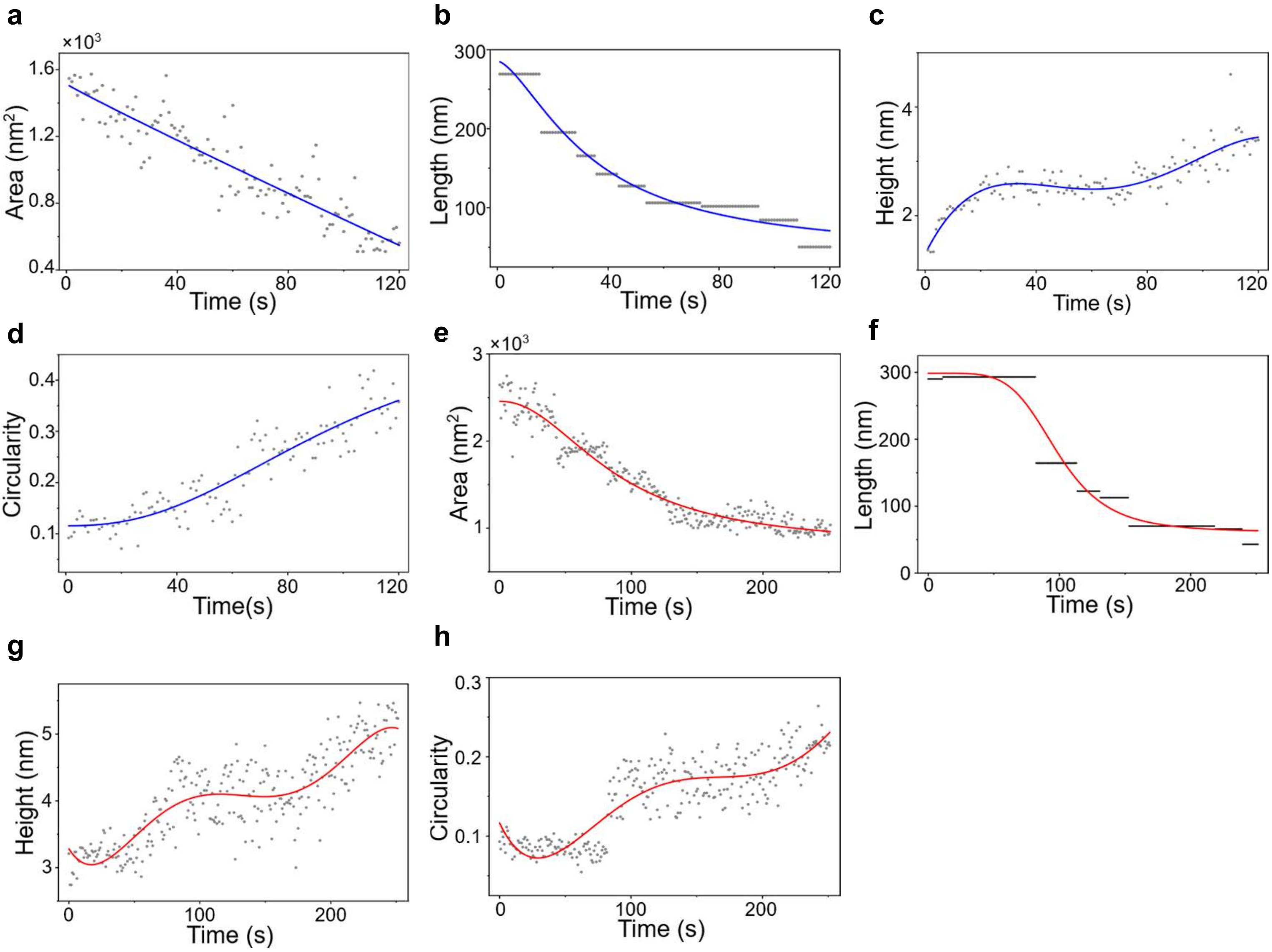


**Figure S4. Physiological properties of single molecule poly A and total RNA.**

**a.b.c.d** Scatter plots showing changes in area, length, height, and circularity of single poly A RNA during scanning in the presence of $Mg^{2+}$. A downward trend was observed for both areas and length where height and circularity increased as RNA formed globular shape. The solid line represents a polynomial fit to the experiment data, illustrating the trend of data during droplet formation.

**e.f.g.h.** Scatter plot showing area, length, height and circularity changes of single total RNA molecules over time during aggregation. The solid line represents a polynomial fit to the experiment data, illustrating the trend of data during aggregates formation.
